# SOLiD-MaP: A Photoproximity Labeling Platform for Small Molecule Binding Site Mapping on RNA

**DOI:** 10.64898/2026.06.16.732620

**Authors:** Lin L. Rietveld, Wei Wu, Filip M. Zawisza, Danny Incarnato, Zeshi Li

## Abstract

RNA-targeting small molecules are emerging as promising therapeutic modalities, but their development requires methods that define binding sites and evaluate RNA target selectivity. Existing approaches for detecting ligand-RNA interactions have provided powerful foundations, yet many rely on direct crosslinking or covalent-capture chemistries whose performance depends on ligand-specific probe design, warhead compatibility, and local reaction geometry. Here, we report Singlet Oxygen footprinting on RNA in a Ligand-Directed manner for Mutational Profiling (SOLiD-MaP), a photoproximity labelling platform for small molecule-RNA interaction analysis. Using the Mango-II aptamer and thiazole orange derivatives as a model system, we establish aniline as an efficient nucleophile for singlet oxygen-mediated RNA labelling and demonstrate target-selective labelling driven by ligand-localized photosensitization. We further show that labelling selectivity can be tuned by chemically constraining the singlet oxygen diffusion with a quencher. Finally, we develop a pairwise reverse transcription stop assay and a mutational profiling with next-generation sequencing readouts to infer ligand-proximal regions and unambiguously map binding sites. SOLiD-MaP provides a new, orthogonal strategy for studying small molecule-RNA recognition and should support RNA-focused mechanism-of-action studies.

---

Small molecule-RNA interactions are emerging as a frontier in chemical biology and drug discovery. Structured RNA elements can be selectively recognized by small molecules to modulate splicing, translation, RNA stability, and other disease-relevant processes.^1-6^ Moreover, transcriptome-scale profiling has revealed that small molecules originally developed for protein targets can also interact extensively with cellular RNAs and, in some cases, alter RNA function.^7^ Defining small molecule-RNA interactions is therefore important not only for developing RNA-targeted therapeutics, but also for understanding off-target pharmacology, anticipating liabilities, and identifying existing chemotypes that may be repurposed as RNA-directed ligands.

Motivated by these opportunities, the field has developed a series of pioneering methods to interrogate RNA-small molecule recognition, including Chem-CLIP,^8,9^ PEARL-seq,^10^ BIVID-MaP,^11^ RBRP^7^ and related covalent-capture strategies (**Figure 1A**).^12-16^ Together, these approaches have established an important methodological foundation for the field and largely operate by converting target engagement into a productive single-event affinity-labeling readout. However, complementary strategies remain of interest because current methods typically require optimization of warhead chemistry and linker architecture, and depend on local reaction geometry. In this regard, photocatalytic proximity labeling methods such as μMap have shown in protein systems that target-localized reactivity can support binding-site mapping without requiring direct crosslink formation.^17,18^ However, comparable proximity labeling-based strategies remain underdeveloped for small molecule-RNA interaction analysis.

**Figure 1.**
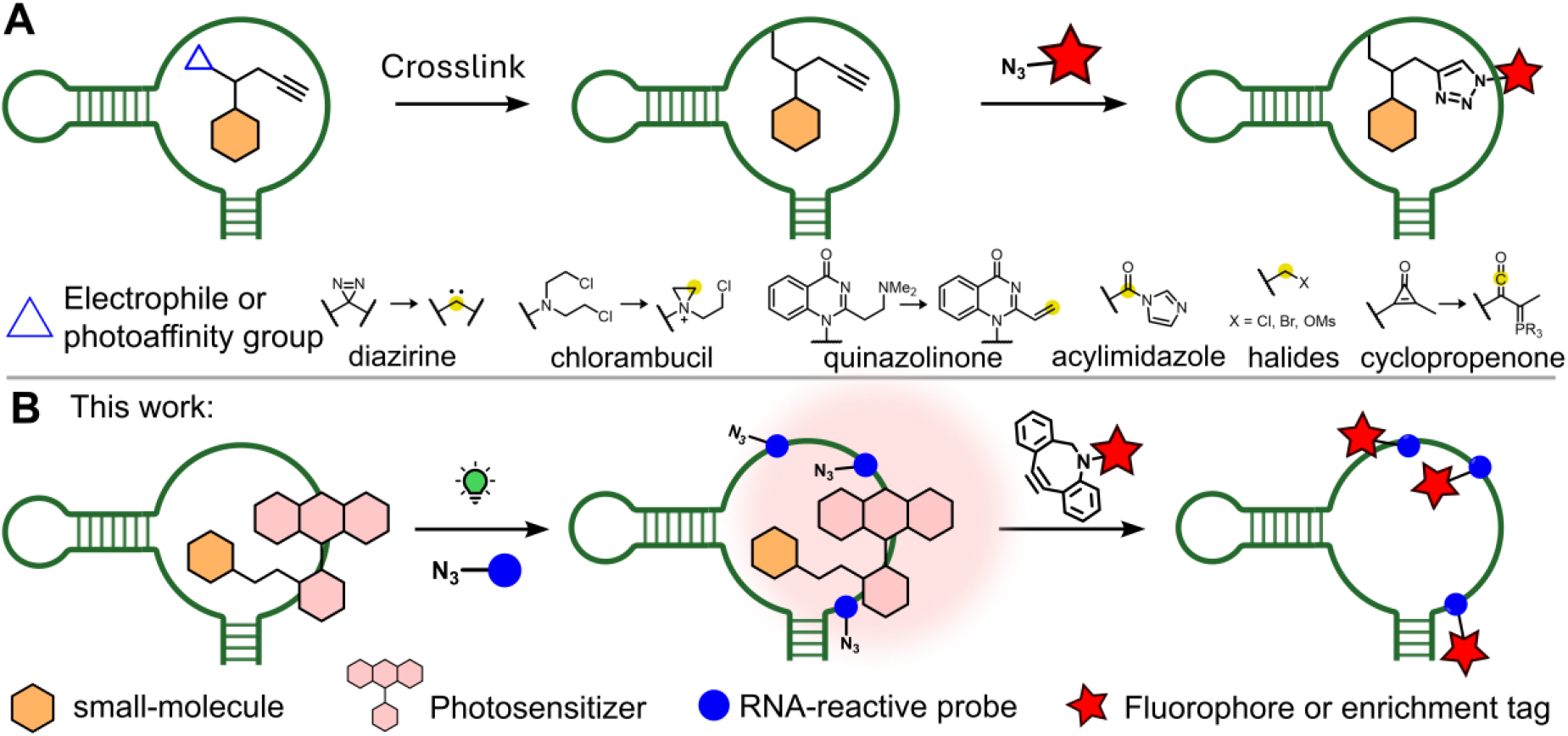
Small molecule-mediated RNA labeling techniques based on the principles of (A) affinity labeling or (B) photoproximity labeling. The electrophilic center is indicated by a yellow solid circle.

Among possible reactive species, singlet oxygen (^1^O_2_) is attractive for RNA proximity labeling because it reacts efficiently with nucleobases and its short lifetime creates a distance-dependent chemical gradient around the photosensitizer. However, ^1^O_2_-mediated RNA labeling has primarily been used to profile broad subcellular RNA populations, where its effective labeling range is well suited to organelle-scale transcriptomics.^19,20^ Recent work has extended this chemistry to probe RNA secondary structures and site-specific labeling of RNA loops.^21^ Binding-region inference within a defined ligand-RNA complex imposes a more stringent requirement: the chemical footprint must be preferentially concentrated near the ligand-proximal region, rather than distributed across the RNA.

Herein, we report Singlet Oxygen footprinting on RNA in a Ligand-Directed manner for Mutational Profiling (SOLiD-MaP), a small molecule–targeted RNA photoproximity labeling platform that leverages ^1^O_2_ reactivity toward nucleobases for binding-site analysis (**Figure 1B**). Using the Mango aptamer and thiazole orange (TO) derivatives as a model system, we show that aniline enables more efficient ^1^O_2_-mediated RNA labeling than alkyl amines, and that the effective labeling range can be chemically constrained with a ^1^O_2_ quencher to improve on-target labeling. We further establish two complementary assays for binding-region inference and higher-resolution site mapping. Together, these results position SOLiD-MaP as an orthogonal approach for studying small molecule-RNA recognition, with potential utility for RNA-focused mechanism-of-action studies in cells.

We first optimized the nucleophile for ^1^O_2_-mediated RNA labeling. Although alkyl amines such as propargylamine are commonly used to trap oxidized G intermediates,^19,20^ we reasoned that aniline may improve labeling efficiency based on their enhanced reactivity toward ^1^O_2_-oxidized histidine in proteins (**Figure 2A**).^22^ Given that the strain promoted click chemistry outperforms copper(II) catalyzed one in RNA conjugation,^23,24^ we compared azide-functionalized alkylamine **APA** and aniline **AnA** using EY-mediated labeling of in vitro-transcribed RNA followed by click chemistry with dibenzocyclooctyne-functionalized Cyanine-5 (DBCO-Cy5) for in-gel fluorescence (**Figure 2B**). Both probes produced light-, EY-, and probe-dependent RNA labeling that was lost after RNase treatment (**Figure 2C, D** and **S1**). Cy5 signal progressively intensified with longer light exposure (**Figure S2**). **AnA** yielded a 3.5-fold higher Cy5 signal than **APA**, establishing aniline as a more efficient nucleophile for ^1^O_2_-mediated RNA labeling (**Figure 2E** and **S1**).

**Figure 2.**
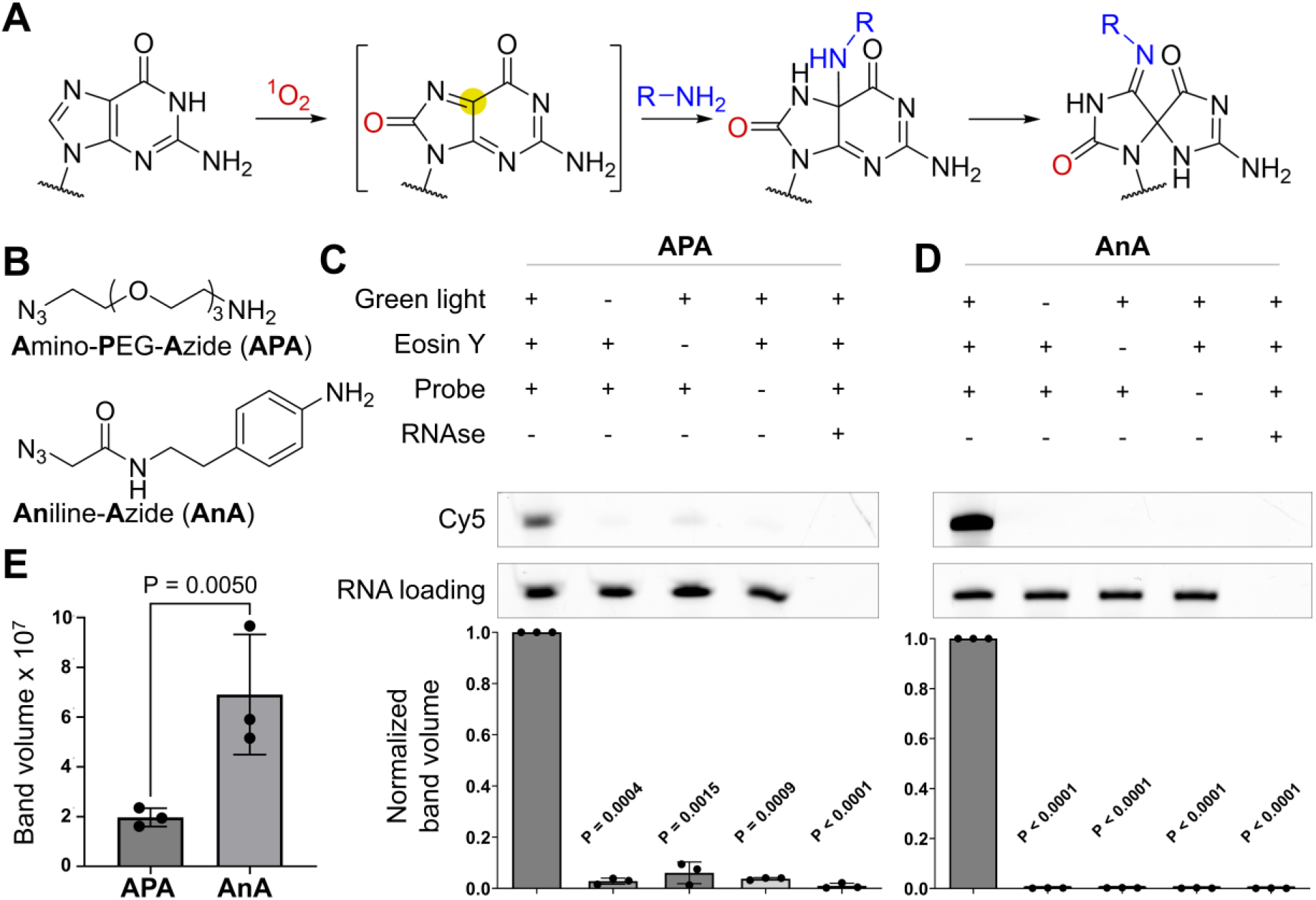
Comparison between alkyl and aryl amines as nucleophiles in singlet oxygen-based RNA labeling. **(A)** Mechanism of singlet oxygen oxidation of guanosine. Yellow solid circle indicates the electrophilic center generated during guanosine oxidation. **(B)** Structures of **APA** and **AnA**. **(C)** and (**D**) Representative in-gel fluorescence of RNA after photochemical labeling of RNA using **APA** (**C**) and **AnA** (**D**). The bar graphs are the quantitative analysis of gel images and represents the mean ± SD of n = 3 independent labelling reactions. (**E**) A side-by-side comparison of the labelling efficiency of **APA** and **AnA**.

Next, attention was focused on establishing a small molecule-based RNA proximity labeling system using Mango-II fluorescence aptamer and the ligand TO as our model system.^25^ Mango binds several TO derivatives and tolerates different spacer modifications.^26^ To conjugate photocatalyst EY to TO, we synthesized **TO-amine** (**Figure 3A**, see Supplementary Information for the synthesis route), which was then reacted with NHS-activated EY to afford the photocatalyst-modified small molecule **TO-EY**. Binding studies using Mango-II RNA and **TO-amine** were performed with fluorescence readout. Strong fluorescence was observed only when **TO-amine** and Mango-II were both present, whereas the fluorescence was substantially reduced either when a control RNA sequence was used, or in the absence of RNA (**Figure 3B**). This confirmed that the triazole-linked TO derivative retains Mango recognition.^27^

**Figure 3.**
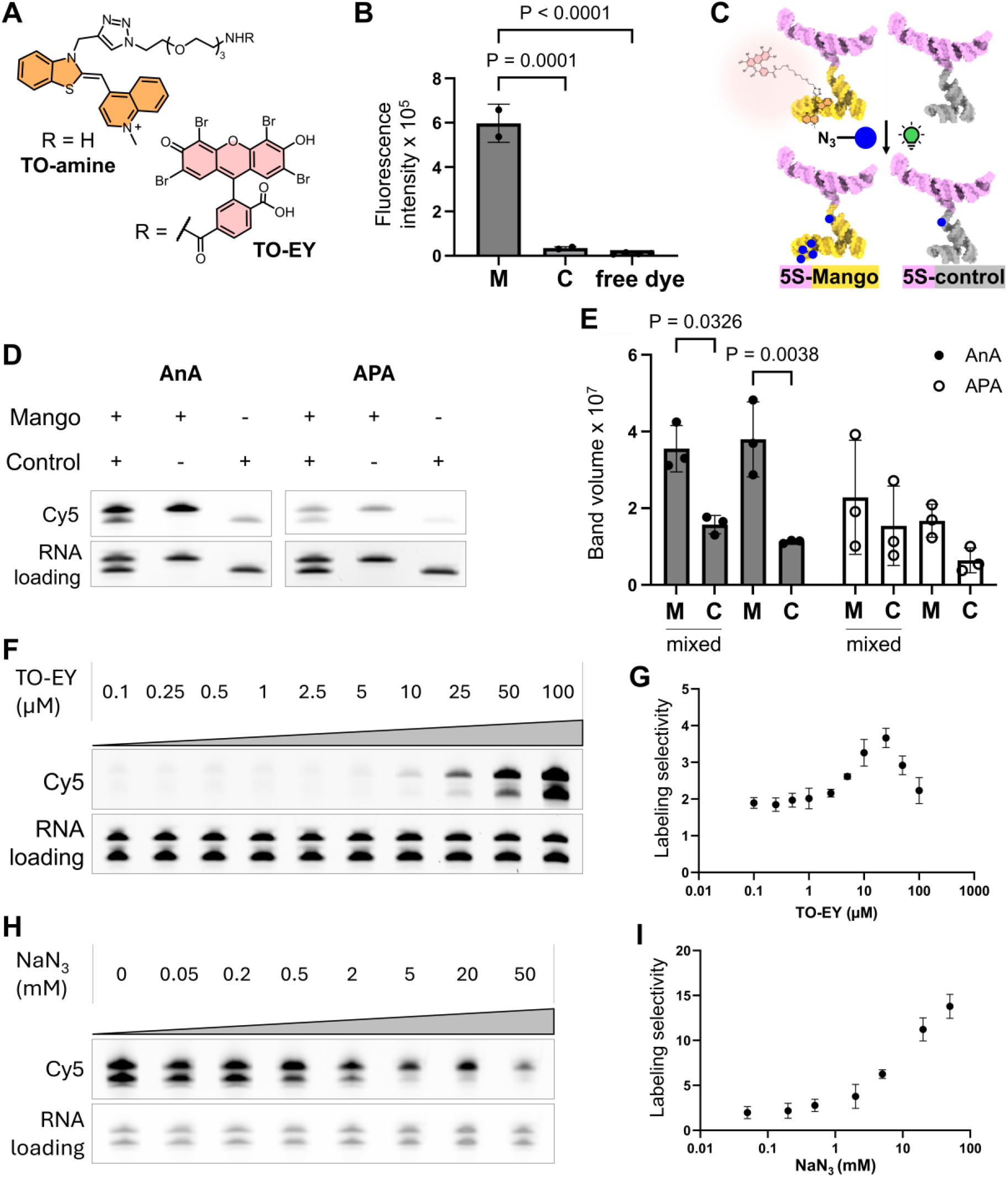
TO-EY guided photoproximity labeling. **(A)** Structures of **TO-amine** and **TO-EY**. **(B)** Fluorescence analysis of **TO-amine** incubated with Mango-II RNA (M), control oligo (C), or with free **TO-amine**. Bar graph represents mean±SD, n = 2 independent replicates. **(C)** Schematic representation of **TO-EY** guided photoproximity labeling of Mango-II (yellow) and control (grey) RNAs constructed with an upstream 5S ribosomal RNA (pink) sequence. Blue solid circle with an azide represents **APA** or **AnA**. **(D)** Representative in-gel fluorescence of RNA after **TO-EY** guided photoproximity labeling using **APA** or **AnA**. 5S-Mango and -control RNAs were either mixed in the same tube or labeled separately in different tubes. **(E)** Quantitative analysis of gel images in panel (**C**). The bar graph represents the mean ± SD of n = 3 independent labelling reactions. M, 5S-mango; C, 5S-control. **(F)** Representative in-gel fluorescence of photoproximity labeled of 5S-Mango and -control using different **TO-EY** concentrations. **AnA** was used as the probe. **(G)** Quantitative analysis of gel images in panel (**E**) regarding the labeling selectivity against **TO-EY** concentration. The dot plot represents the mean ± SD of n = 3 independent labelling reactions. **(H)** Representative in-gel fluorescence of photoproximity labeled RNAs in the presence of different amounts of NaN_3_. **(I)** Quantitative analysis of gel images in panel (**G**) regarding the labeling selectivity against NaN_3_ concentration. The dot plot represents the mean ± SD of n = 3 independent labelling reactions.

In the **TO-EY** proximity labeling system, photosensitizer recruitment to Mango-II should preferentially label the target RNA over a non-binding control (**Figure 3C**). We tested this in a same-tube labeling assay using *in vitro* transcribed 5S ribosomal RNA-derived constructs: a 212-nt 5S-F30-Mango-II target and a 199-nt control in which Mango-II was replaced by four nucleotides, GAAA, with the remaining sequence unchanged. We observed that the **AnA** probe labels the 5S-Mango RNA significantly more than the control (**Figure 3D** and **E**). Consistent results were obtained when the labeling was performed separately for the two RNAs. Unlike **AnA**, the **APA** probe showed a non-significant difference between target and control labeling. Blue light irradiation had a similar result (**Figure S3**). A loss of selectivity was observed when **TO-amine** was used to compete for Mango-II binding (**Figure S4**). To confirm the proximity effect, a 40-nt sequence (designated 5S binder) was hybridized to a shared region in 5S-Mango and 5S-control was used. Upon separate labeling of 5S binder-hybridized 5S-Mango or 5S-control using **TO-EY**, urea-PAGE showed markedly stronger Cy5 labeling of the binder hybridized to 5S-Mango, indicating that enhanced labeling primarily reflects closer EY proximity rather than sequence composition (**Figure S5**). Our results demonstrate that small molecule-directed target-selective labeling can be achieved with the **AnA** probe, which was used for the rest of the study.

We next asked how excess ligand-photosensitizer conjugate affects target selectivity. Selectivity was defined as the Cy5 band-volume ratio of target RNA to control RNA after separate normalization to RNA loading by ethidium bromide (EtBr) staining. As **TO-EY** concentration increased from 0.1 to 100 µM, Cy5 signals intensified for both RNAs, while target selectivity peaked at 25 µM with an approximate target/control ratio of 3.7 before declining at higher concentrations (**Figure 3F** and **G**). Importantly, labeling remained 5S-Mango-selective even under excess **TO-EY**, indicating that target preference is retained despite the presence of largely unbound photosensitizers.

Because the transiently reactive species in our system is ^1^O_2_, we hypothesized that inclusion of a ^1^O_2_ quencher, such as sodium azide (NaN_3_),^28^ would constrain its diffusion and thereby suppress off-target labeling. Indeed, increasing NaN_3_ concentration progressively enhanced target RNA selectivity (**Figure 3H** and **I**). While Cy5 intensity decreased for both target and control RNA, the decline was markedly steeper for the control RNA as quencher concentration increased. These findings establish that reactive-species quenching can be leveraged to tune the effective labeling radius, enabling a controllable trade-off between labeling efficiency and target selectivity.

We next asked whether photoproximity labeling could provide binding-site information on the RNA target by developing a pairwise reverse transcription (RT) stop assay combined with real-time quantitative polymerase chain reaction (qPCR). Each 5S-Mango RNA sample was split in two and reverse-transcribed with primers upstream or downstream of Mango-II, followed by qPCR using primers F-ex/R-ex to quantify the Mango-II-excluding RT product (Mango-upstream RT), and F-in/R-in to quantify the Mango-II-containing region (downstream RT) (**Figure 4A**). Unlabeled RNA gave comparable apparent quantities for both RT products (**Figure 4B**), whereas **TO-EY**-directed labeling (in the presence of 10 mM NaN_3_) selectively reduced the Mango-II-containing RT product, consistent with local **AnA** modification causing RT stops that prevent F-in/R-in amplification. By contrast, free EY labeling reduced both RT products similarly, indicating more evenly distributed RT stops across the 5S and Mango-II regions.

**Figure 4.**
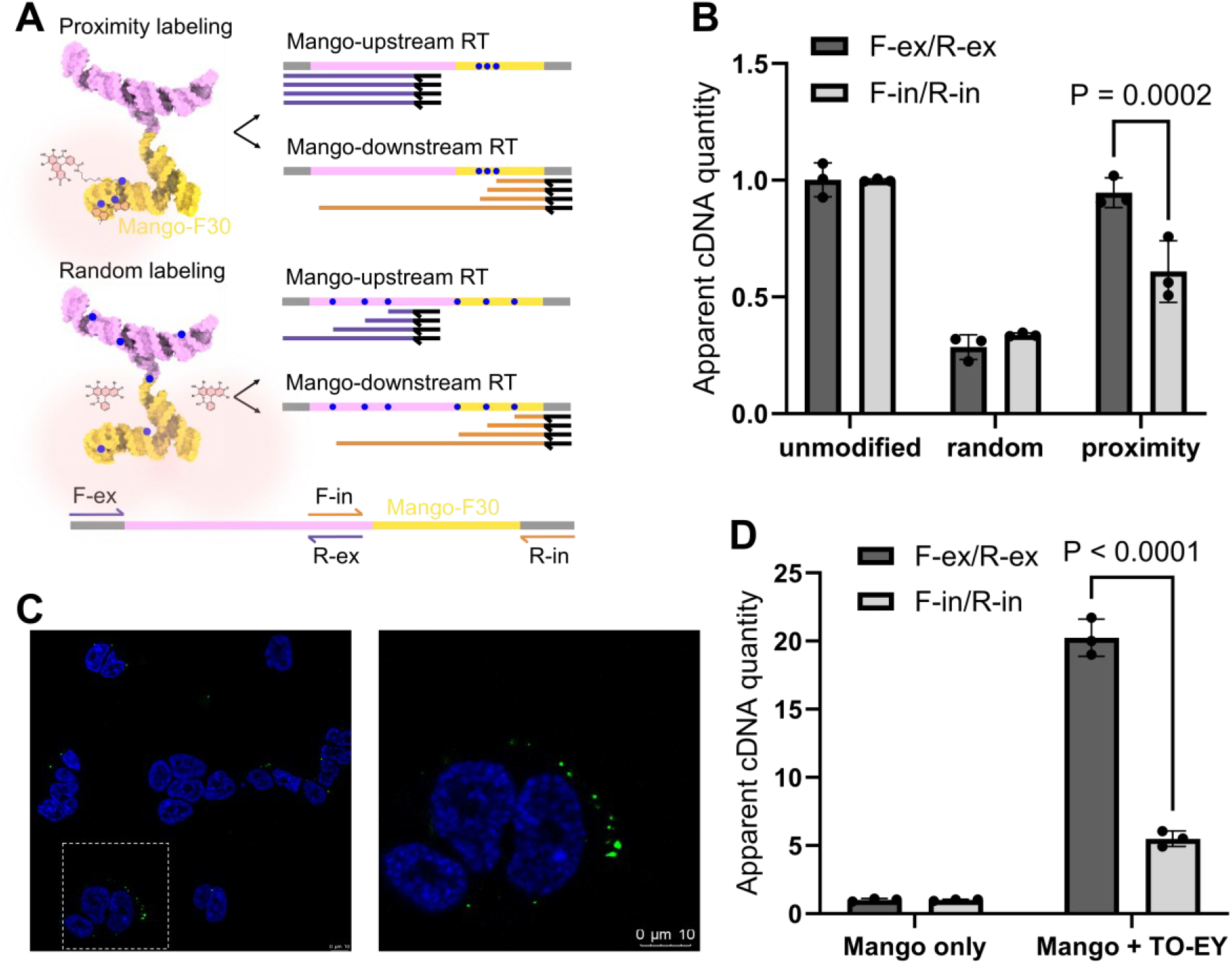
RT stop combined with qPCR assay for binding regio inference. **(A)** Schematic representation of samples. Black single headed arrow, RT primer; purple and orange lines, cDNA. **(B)** Relative quantities of PCR-amplifiable RT product in unmodified, randomly labeled and proximity labeled 5S-Mango, performed in vitro. Bar graph represents mean ± SD. n = 3 technical replicates. See **Figure S6** for the other independent replicate. See supplementary information for calculation. **(C)** Confocal imaging confirmed precomplexed 5S-Mango and **TO-amine** stayed intact after transfection. n = 2 independent cell cultures. Left, broader field of view. Right, zoomed-in view of the region defined by a dashed frame. Scale bar, 10 µm. **(D)** Relative quantities of PCR-amplifiable RT product of proximity labeled 5S-Mango in living cells. Bar graph represents mean±SD. See **Figure S6** for the other independent replicate.

Next, we sought to investigate whether the binding site inference can be achieved in a living-cell context. We first confirmed the preformed TO-Mango complex can stay intact after transfection, as specific fluorescence signals were detectable with **TO-amine** precomplexed with 5S-Mango (**Figure 4C** and **S6**). To perform photoproximity labeling in living cells, precomplexed **TO-EY** and 5S-Mango were transfected into HEK293T cells, after which light irradiation was performed in the presence of a biotin-modified aniline probe (see Supplementary information for structure and synthesis). The total RNA post-labeling was extracted and subjected to streptavidin-bead pull-down to enrich labeled RNA. Consistent within vitro labeling, we observed a significant reduction in apparent quantity of Mango-downstream RT product, relative to the upstream one in **TO-EY** in cell proximity labeling (**Figure 4D**). Taken together, the pairwise RT stop detection with real-time qPCR allows for inferring the location of the small molecule binding site on an RNA both *in vitro* and in a cellular context.

Motivated by the above results, we then sought to unambiguously map the small molecule interacting region using Next-Generation sequencing (NGS). The photochemically labeled RNAs were subjected to RT under mutational profiling (MaP) condition,^29^ which forces the RT enzyme to read through the modified bases and introduce mutations in the cDNA around those positions (**Figure 5A**).

**Figure 5.**
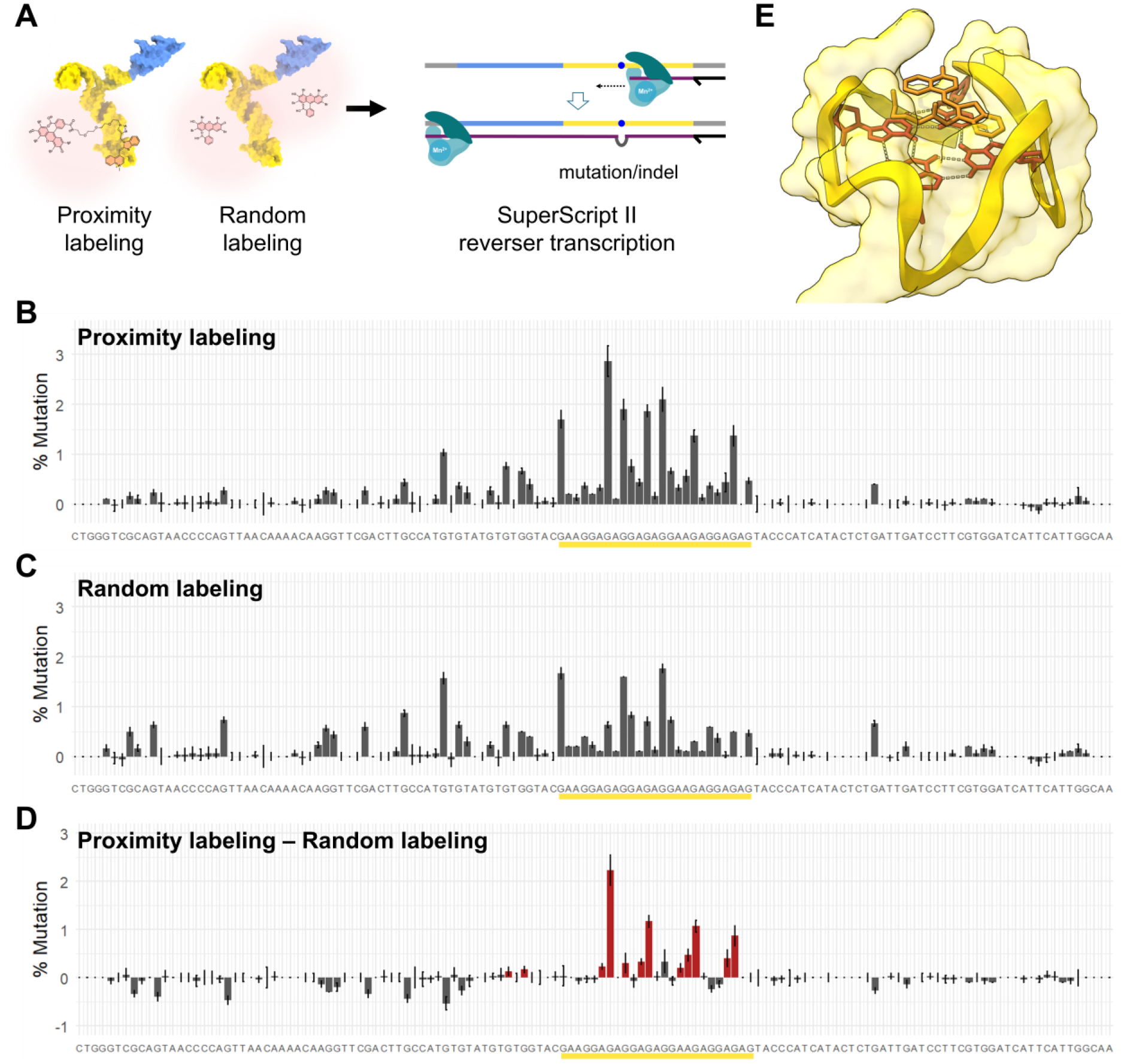
Mutational profiling of proximity and randomly labeled RNA. **(A)** Schematic representation of experimental procedures. The singlet oxygen quencher used in the experiment, NaN_3_ (5 mM), is not displayed. **(B)** and (**C**) Bar graphs (mean ± SD, n = 3) of mutation rates of RNA proximity labeled with **TO-EY** (**B**) or randomly labeled with free EY (**C**). The yellow underscore indicates the Mango II sequence. The mutation rates at each position was subtracted by that of unmodified 5S-mango. **(D)** Mutation rate difference obtained by subtracting (**B**) with (**C**). Red coloring indicates position with adjusted p < 0.05 AND log_2_(fold change) > 1 (limma). **(E)** Mango II (yellow backbone and surface) in complex with TO1-biotin ligand (orange sticks) (PDB ID 6C63). Residues with mutation rate > 0.5% were shown as red sticks.

When the Mango-II-containing RNA were proximity labeled with **TO-EY** in the presence of 5 mM NaN_3_, we observed that highly mutated positions were clustered at the Mango-II region, while the rest harbors much lower mutation rates after background subtraction (**Figure 5B**, see **Figure S7** for unprocessed mutation rates). Switching **TO-EY** to free EY afforded a more scattered mutation rate distribution across the RNA sequence (**Figure 5C**). This random labeling reflects the intrinsic reactivities of different Gs towards ^1^O_2_-mediated oxidation and **AnA** modification.^21^ Omitting the photosensitizer produced little detectable mutation along the entire sequence after background subtraction (**Figure S7F and G**). Remarkably, subtracting **TO-EY** RNA mutation rate with that of free EY, we observed four Gs within the Mango-II region with particularly larger mutation rate difference than the rest. These G residues are exactly those constituting the G quartet directly interacting with the TO ligand (**Figure 5E**). In contrast, the Gs of the bottom two rG4 planes had much lower to no difference in mutation rates, despite the proximity to the binding site. Taken together, the small molecule-directed ^1^O_2_ footprinting can thus unambiguously define the binding site on RNA.

In summary, we introduce SOLiD-MaP, a small-molecule photoproximity labeling strategy for RNA binding-site analysis, and establish the platform comprehensively in a model system, from probe design and target-selective labeling to binding-site inference. Three features underpin SOLiD-MaP. First, a key chemical advance is the identification of aniline as a more efficient and selective nucleophilic probe than alkyl amines for ^1^O_2_-mediated RNA labeling. Second, addition of the quencher NaN_3_ reduces the effective labeling range and improves target selectivity. Third, the method harnesses the high efficiency of ^1^O_2_-mediated chemistry without requiring probe reactivity to be restricted to the binding pocket, as in μMap. Rather, binding-site information arises from elevated ^1^O_2_ reactivity at specific G residues near the bound ligand-photosensitizer, compared to the intrinsic reactivity profile (generated by ramdom labeling using free EY). Because the enhanced G reactivity is governed primarily by proximity to the catalytic center, SOLiD-MaP should be less dependent on linker architecture than electrophile-or photoaffinity-based methods, in which precise placement of the correct reactive warhead is often essential. Together, these results establish a complementary, orthogonal alternative to direct (photo)crosslinking for studying small molecule-RNA interactions and identifying RNA binding sites.

## Supporting information

Integrated Supplementary Information

## References

1 Warner, K. D., Hajdin, C. E. & Weeks, K. M. Principles for targeting RNA with drug-like small molecules. Nat Rev Drug Discov 17, 547–558 (2018). 10.1038/nrd.2018.93

2 Wang, J., Schultz, P. G. & Johnson, K. A. Mechanistic studies of a small-molecule modulator of SMN2 splicing. Proc Natl Acad Sci U S A 115, E4604–E4612 (2018). 10.1073/pnas.1800260115

3 Childs-Disney, J. L., Yang, X., Gibaut, Q. M. R. et al. Targeting RNA structures with small molecules. Nat Rev Drug Discov 21, 736–762 (2022). 10.1038/s41573-022-00521-4

4 Aguilar, R., Spencer, K. B., Kesner, B. et al. Targeting Xist with compounds that disrupt RNA structure and X inactivation. Nature 604, 160–166 (2022). 10.1038/s41586-022-04537-z

5 Tong, Y., Lee, Y., Liu, X. et al. Programming inactive RNA-binding small molecules into bioactive degraders. Nature 618, 169–179 (2023). 10.1038/s41586-023-06091-8

6 Zhang, C., Borovska, I., Iobashvili, T. et al. RNA functional modulation by Mitoxantrone via RNA structural ensemble repartitioning. Nat Commun 17 (2026). 10.1038/s41467-026-70801-9

7 Fang, L., Velema, W. A., Lee, Y. et al. Pervasive transcriptome interactions of protein-targeted drugs. Nat Chem 15, 1374–1383 (2023). 10.1038/s41557-023-01309-8

8 Guan, L. & Disney, M. D. Covalent small-molecule-RNA complex formation enables cellular profiling of small-molecule-RNA interactions. Angew Chem Int Ed Engl 52, 10010–10013 (2013). 10.1002/anie.201301639

9 Yang, W. Y., Wilson, H. D., Velagapudi, S. P. & Disney, M. D. Inhibition of Non-ATG Translational Events in Cells via Covalent Small Molecules Targeting RNA. J Am Chem Soc 137, 5336–5345 (2015). 10.1021/ja507448y

10 Mukherjee, H., Blain, J. C., Vandivier, L. E. et al. PEARL-seq: A Photoaffinity Platform for the Analysis of Small Molecule-RNA Interactions. ACS Chem Biol 15, 2374–2381 (2020). 10.1021/acschembio.0c00357

11 Miyashita, E., Onizuka, K., Chen, Y. et al. Systematic identification of variant-specific RNA structure-small molecule interactions exemplified by RNA G-quadruplexes. Nat Commun 17 (2026). 10.1038/s41467-026-70097-9

12 Bereiter, R., Flemmich, L., Nykiel, K. et al. Engineering covalent small molecule-RNA complexes in living cells. Nat Chem Biol 21, 843–854 (2025). 10.1038/s41589-024-01801-3

13 Chen, S., Sibley, C. D., Latifi, B. et al. Bioorthogonal Cyclopropenones for Investigating RNA Structure. ACS Chem Biol 19, 2406–2411 (2024). 10.1021/acschembio.4c00633

14 Balaratnam, S., Rhodes, C., Bume, D. D. et al. A chemical probe based on the PreQ(1) metabolite enables transcriptome-wide mapping of binding sites. Nat Commun 12, 5856 (2021). 10.1038/s41467-021-25973-x

15 Crielaard, S., Maassen, R., Vosman, T., Rempkens, I. & Velema, W. A. Affinity-Based Profiling of the Flavin Mononucleotide Riboswitch. J Am Chem Soc 144, 10462–10470 (2022). 10.1021/jacs.2c02685

16 Yesley, P., Poulladofonou, G., Incarnato, D. & Velema, W. A. Site-Selective Ligand Selection by Mutational Profiling for Covalent RNA Targeting. Angew Chem Int Ed Engl 65, e17243 (2026). 10.1002/anie.202517243

17 Trowbridge, A. D., Seath, C. P., Rodriguez-Rivera, F. P. et al. Small molecule photocatalysis enables drug target identification via energy transfer. Proc Natl Acad Sci U S A 119, e2208077119 (2022). 10.1073/pnas.2208077119

18 Huth, S. W., Oakley, J. V., Seath, C. P. et al. muMap Photoproximity Labeling Enables Small Molecule Binding Site Mapping. J Am Chem Soc 145, 16289–16296 (2023). 10.1021/jacs.3c03325

19 Li, Y., Aggarwal, M. B., Nguyen, K., Ke, K. & Spitale, R. C. Assaying RNA Localization in Situ with Spatially Restricted Nucleobase Oxidation. ACS Chem Biol 12, 2709–2714 (2017). 10.1021/acschembio.7b00519

20 Wang, P., Tang, W., Li, Z. et al. Mapping spatial transcriptome with light-activated proximity-dependent RNA labeling. Nat Chem Biol 15, 1110–1119 (2019). 10.1038/s41589-019-0368-5

21 Shentu, J., Jiang, Z., Tan, Q. & Fang, L. Structure-Encoded Oxidation Enables Nucleotide-Resolved RNA Editing, Conjugation, and Structural Probing. Angew Chem Int Ed Engl, e1434064 (2026). 10.1002/anie.1434064

22 Zhai, Y., Zhang, X., Chen, Z. et al. Global profiling of functional histidines in live cells using small-molecule photosensitizer and chemical probe relay labelling. Nat Chem 16, 1546–1557 (2024). 10.1038/s41557-024-01545-6

23 Hu, H., Flynn, N., Zhang, H. et al. SPAAC-NAD-seq, a sensitive and accurate method to profile NAD(+)-capped transcripts. Proc Natl Acad Sci U S A 118 (2021). 10.1073/pnas.2025595118

24 Liang, J., Jia, H., Li, L., Li, X. & Li, Y. beta-Difluoroalkylamine as a Motif for Singlet Oxygen-Mediated Proximity Labeling in Living Cells. Org Lett 23, 4640–4644 (2021). 10.1021/acs.orglett.1c01377

25 Autour, A., Jeng, S. C. Y., Cawte, A. D. et al. Fluorogenic RNA Mango aptamers for imaging small non-coding RNAs in mammalian cells. Nat Commun 9, 656 (2018). 10.1038/s41467-018-02993-8

26 Trachman, R. J., 3rd, Abdolahzadeh, A., Andreoni, A. et al. Crystal Structures of the Mango-II RNA Aptamer Reveal Heterogeneous Fluorophore Binding and Guide Engineering of Variants with Improved Selectivity and Brightness. Biochemistry 57, 3544–3548 (2018). 10.1021/acs.biochem.8b00399

27 Bychenko, O. S., Khrulev, A. A., Svetlova, J. I. et al. Red light-emitting short Mango-based system enables tracking a mycobacterial small noncoding RNA in infected macrophages. Nucleic Acids Res 51, 2586–2601 (2023). 10.1093/nar/gkad100

28 Qian, W., Kumar, N., Roginskaya, V. et al. Chemoptogenetic damage to mitochondria causes rapid telomere dysfunction. Proc Natl Acad Sci U S A 116, 18435–18444 (2019). 10.1073/pnas.1910574116

29 Siegfried, N. A., Busan, S., Rice, G. M., Nelson, J. A. & Weeks, K. M. RNA motif discovery by SHAPE and mutational profiling (SHAPE-MaP). Nat Methods 11, 959–965 (2014). 10.1038/nmeth.3029

