## Supplementary material for "SOLiD-MaP: A Photoproximity Labeling Platform for Small Molecule Binding Site Mapping on RNA": Integrated Supplementary Information

**Supplementary Figures S1 – S7** Page S1

**Experimental Section** Page S9

1. RNA and DNA sequences used in this study
2. Custom light chamber setup
3. In vitro transcription
4. General procedures for RNA labeling, purification and analysis
5. Experiment-specific labeling procedures
6. Other procedures
7. Synthesis procedures

**Supplementary References** Page S28

#### Supplementary Figures

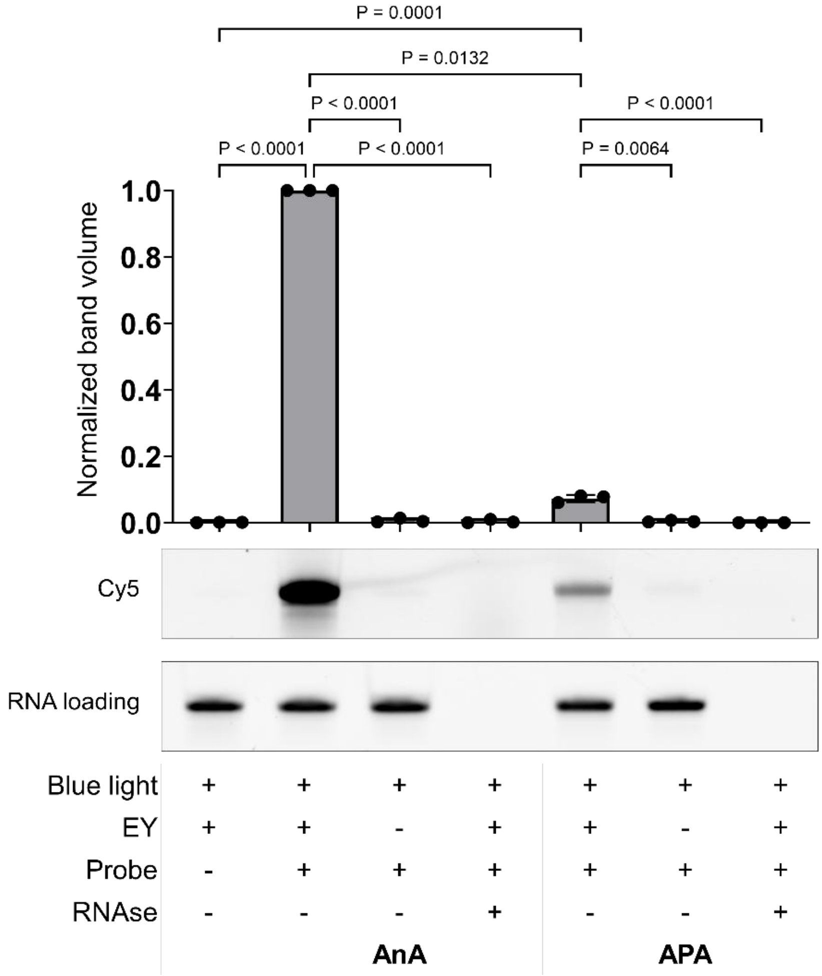


**Figure S1. RNA labeling under blue light activation using free EY with AnA or APA as nucleophiles**. Cy5 band volumes are adjusted to the corresponding RNA loading signal (EtBr signal) and normalized to the highest-intensity band within each gel. Data represent mean ±SD from n = 3 independent labeling reactions. Data were log-transformed and analyzed by one-way ANOVA, followed by Šidák's multiple-comparisons correction.

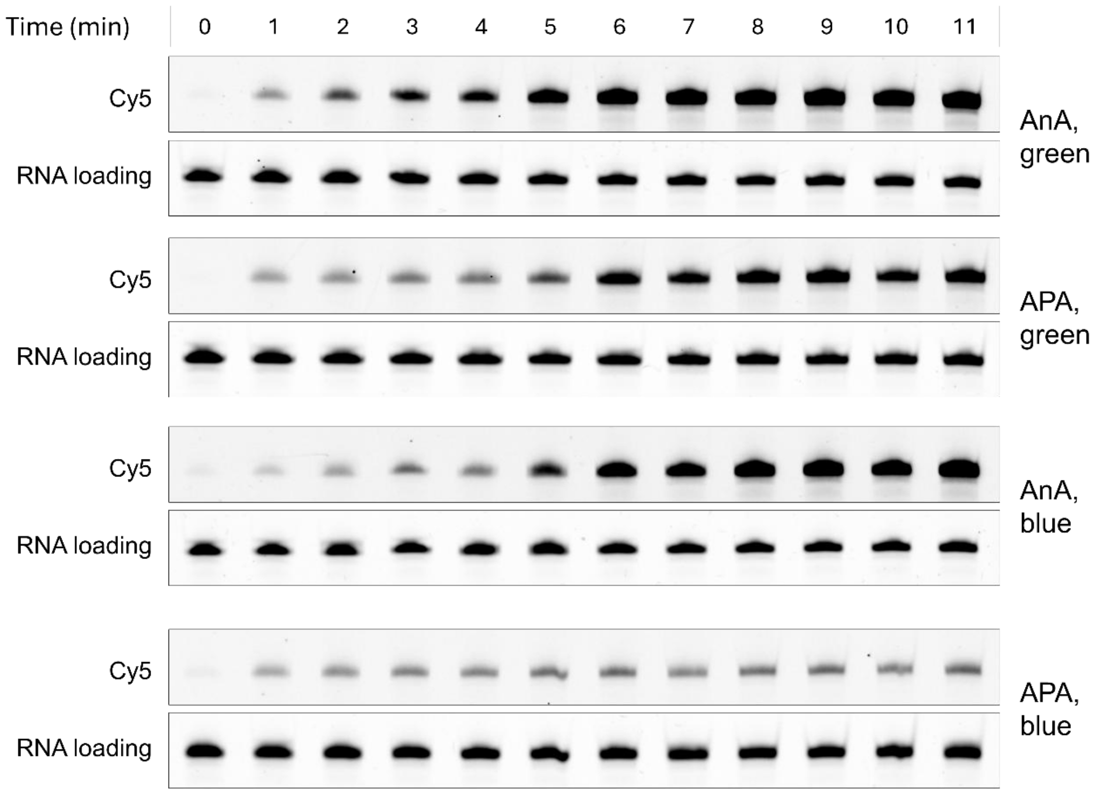


**Figure S2. Time-dependent RNA labeling under continuous irradiation**. RNA labeling using free EY was irradiated under either green or blue light and collected individually at 1-minute intervals.

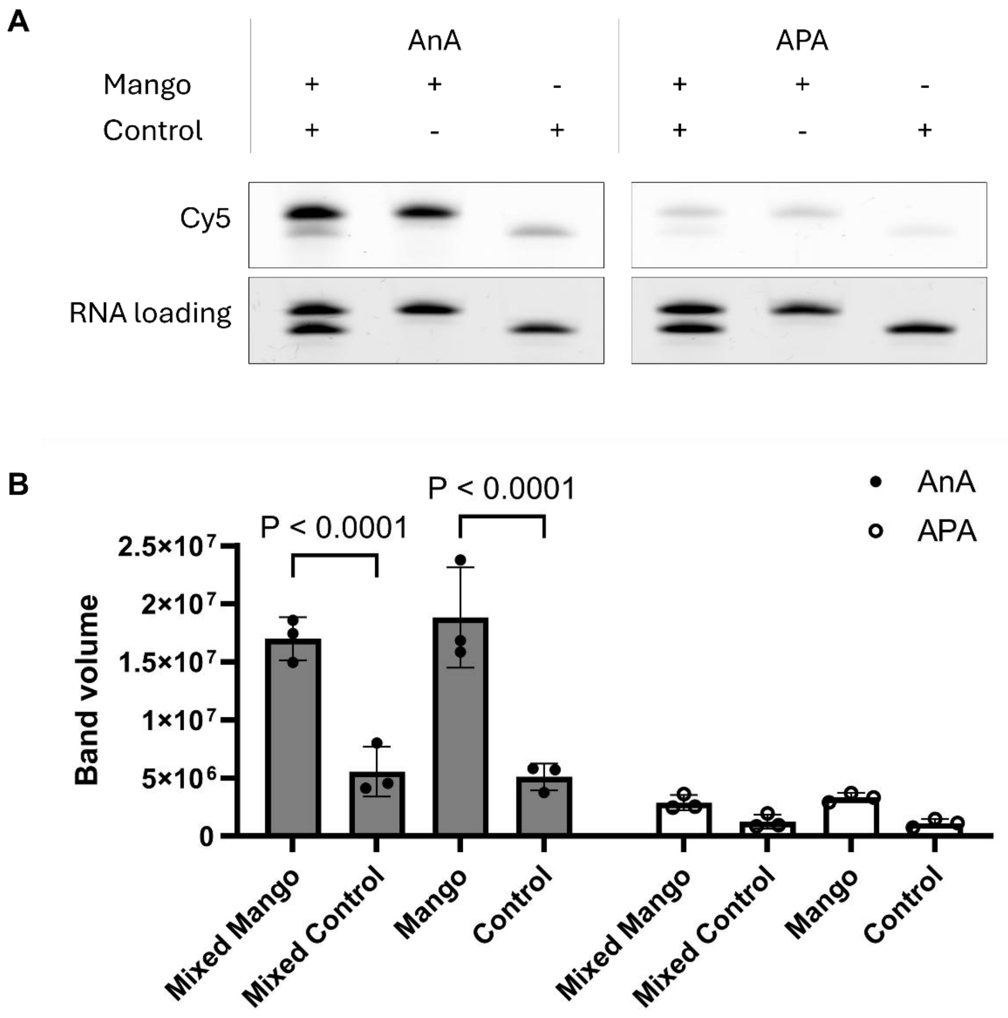


**Figure S3. Blue light activated RNA labeling directed by TO-EY**. (**A**) Representative in-gel fluorescence of RNA after TO-EY guided blue-light photoproximity labeling using **APA** or **AnA**. 5S-mango and -control RNAs were either mixed in the same tube (first lane) or labeled separately in different tubes (second and third lane). (**B**) Quantitative analysis of gel images in panel (**A**). Cy5 signals in gel were quantified with their band volumes, corrected by RNA loading (band volume in EtBr channel). The bar graph represents the mean ± SD of n = 3 independent labelling reactions and were analyzed by two-way ANOVA followed by Tukey's multiple-comparisons test.

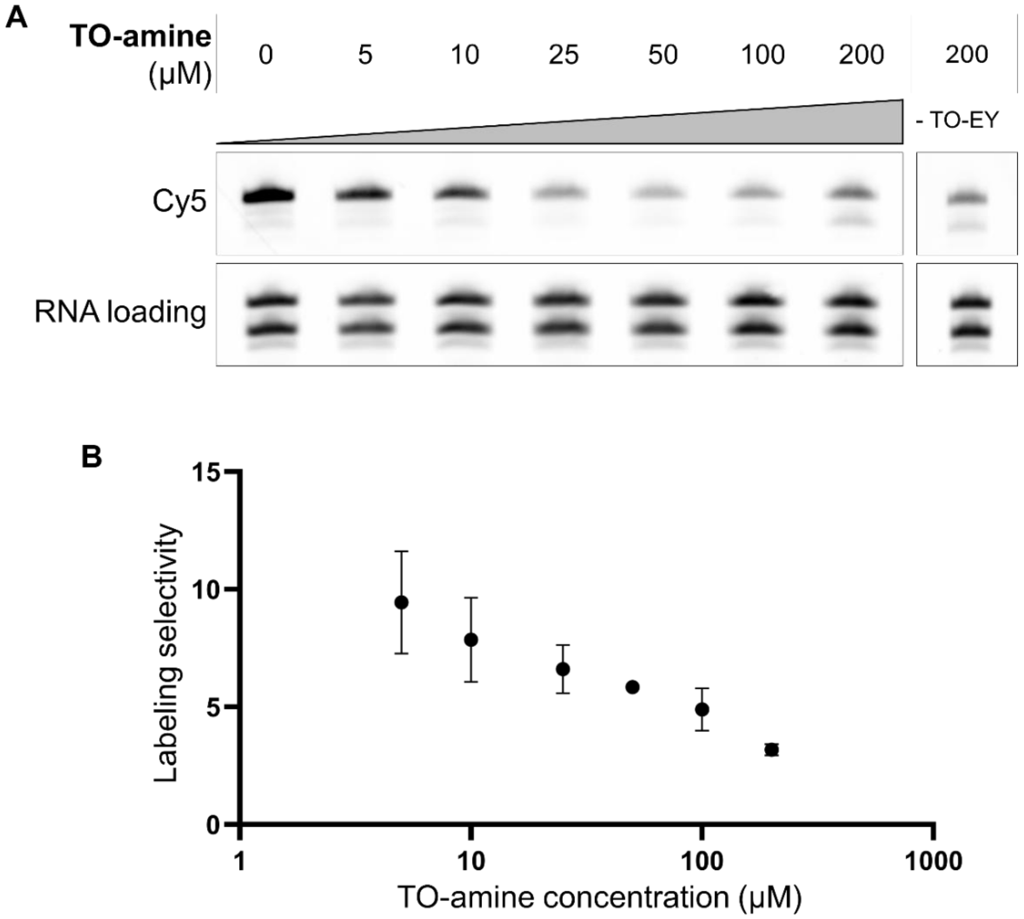


**Figure S4. Effect of competitive binding on TO-EY-directed labeling selectivity**. (A) Representative in-gel fluorescence of RNA after TO-EY (10 μM) guided photoproximity labeling in the presence of increasing concentrations of the competitor TO-amine (5-200 μM). (B) ) Quantitative analysis of gel images in panel (**A**). Scatter plot represents mean ±SD of RNA loading-adjusted Mango/control Cy5 band volume ratios plotted against TO-amine concentration. n = 2 independent labeling experiments. Cy5 bands given rise to by TO-amine alone indicates TO itself is a weak singlet oxygen generator.

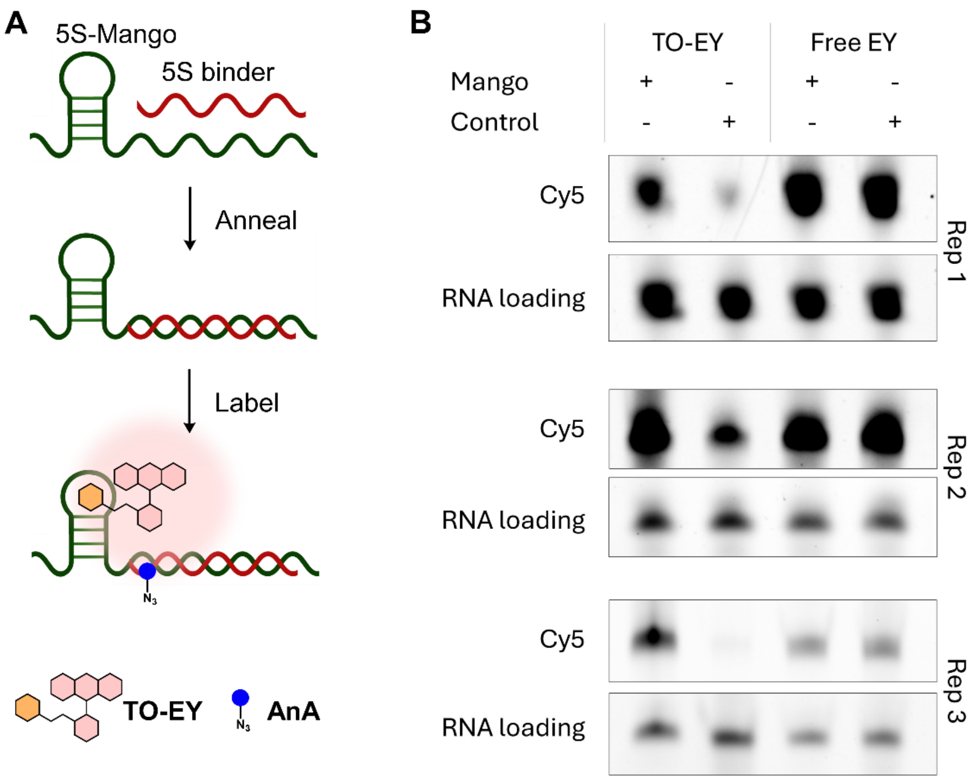


**Figure S5. TO-EY-directed proximity labeling and free-EY random labeling of 5S binder.** (**A**) Schematic representation for proximity-dependent labeling of 5S-binder RNA annealed to 5S-Mango RNA. (**B**) Urea-PAGE analysis of 5S-binder RNA. 5S binder was annealed to either 5S-Mango or 5S-control RNA and labeled using either TO-EY or free EY with **AnA** as the nucleophile. In-gel fluorescence analysis demonstrates enhanced TO-EY-directed labeling of 5S-binder hybridized to 5S-Mango relative to hybridized to 5S-control RNA. No selective labeling was observed under free EY random labeling. Gels from n = 3 independent experiments are shown.

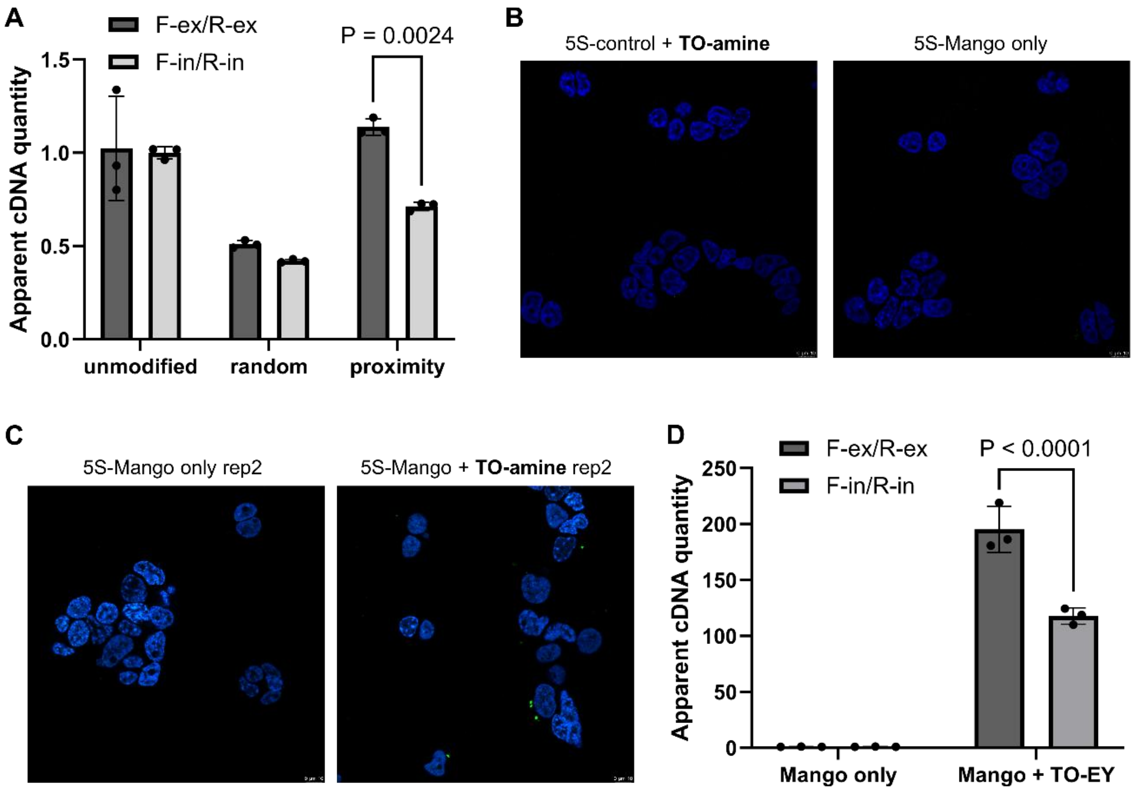


**Figure S6. Additional data of in-vitro and in-cell pair-wise RT stop qPCR assays**. (**A**) Relative quantities of PCR-amplifiable RT product in unmodified, randomly labeled and proximity labeled 5S-Mango, performed in vitro, as an independent replicate of **Figure 4B**. Bar graph represents mean ± SD. n = 3 technical replicates. (**B**) Negative controls in confocal imaging in **Figure 4C**. (**C**) Confirmation that precomplexed 5S-Mango and **TO-amine** stayed intact after transfection, as independent experiment of **Figure 4C**. Scale bar, 10 µm. (**D**) Relative quantities of PCR-amplifiable RT product of proximity labeled 5S-Mango in living cells, as an independent replicate of **Figure 4D**. Bar graph represents mean±SD.

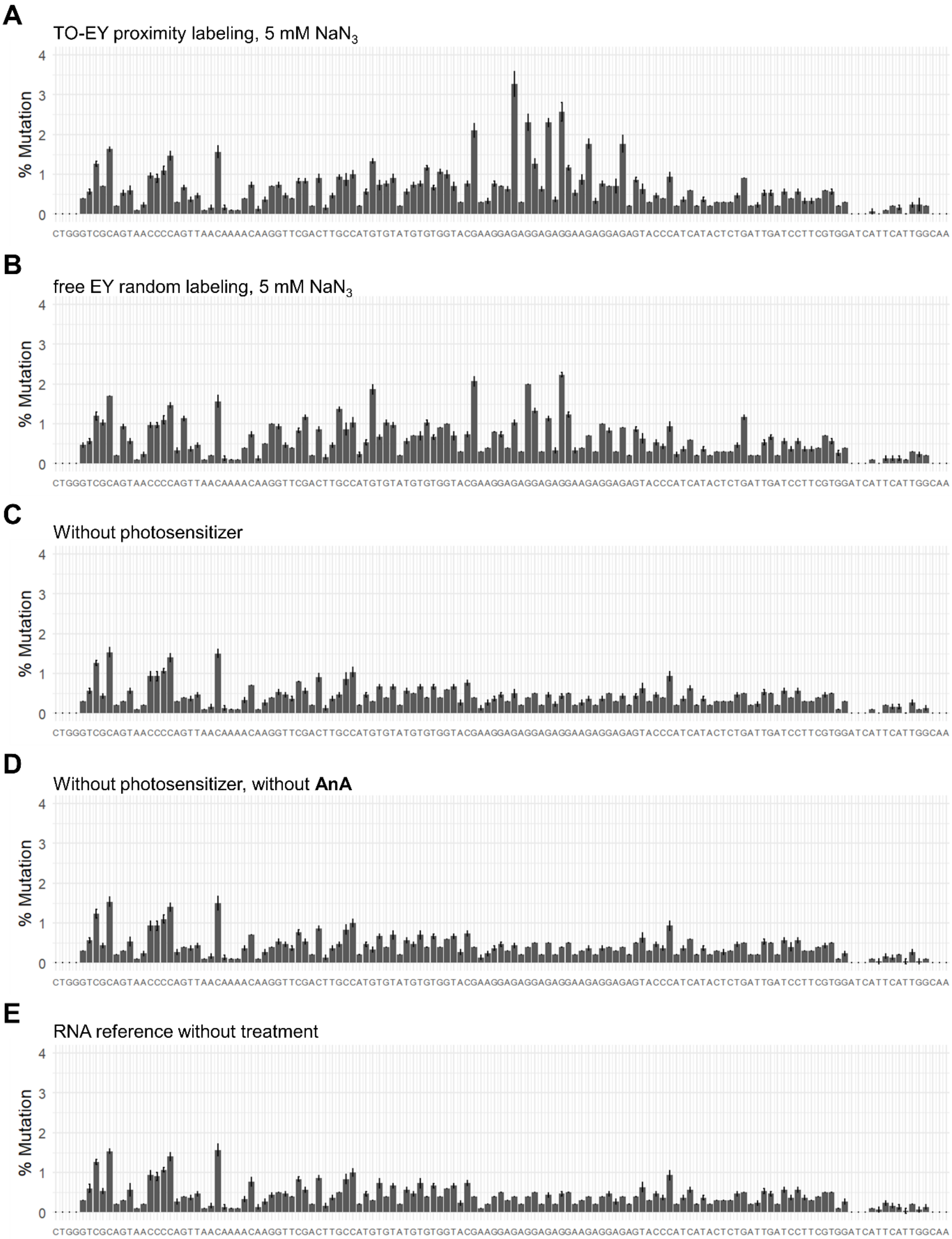


Figure continues on next page

F

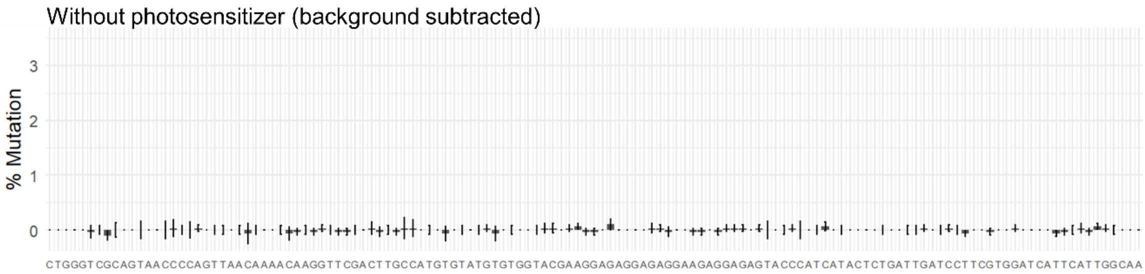


G

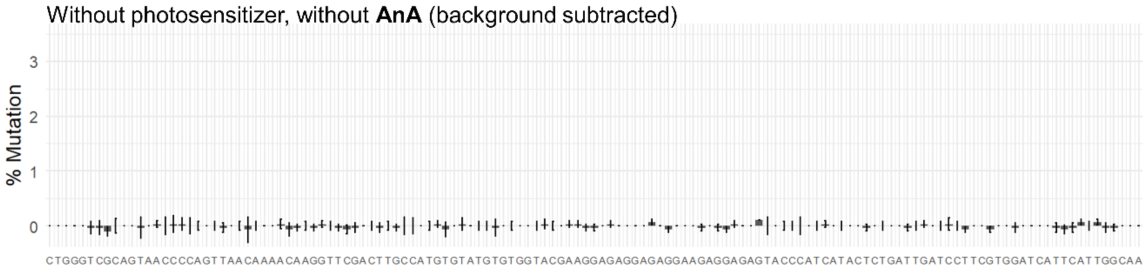


**Figure S7. Mutation profiles of RNA samples without background subtraction**. The untreated RNA reference (**E**) served as background for (**A**) – (**D**). For all samples, bar graphs (mean ±SD) represent n = 3 independent labeling reactions.

### Experimental Section

#### 1. RNA and DNA sequences used in this study

Sequence IDs are called out in the procedures.

##### 1.1 RNA sequences

| Name | ID | Sequence |
|---|---|---|
| Mango-II | LR01 | GCACGUACGA AGGAGAGGAG AGGAAGAGGA GAGUACGUGC |
| Control oligo | LR02 | GCGAAAGCUG GGGAGC |
| 5S-Mango | LR03 | GGUCUACGGC CAUACCACCC UGAACGCGCC CGAUCUCGUC UGAUCUCGGA AGCUAAGCAG GGUCGGGCCU GGUUAGUACU UGGAUGGGAG ACCGCCUGGG AAUACCGGGU GCUGUAGGCG UCGACUUGCC AUGUGUAUGU GGGUACGAAG GAGAGGAGAG GAAGAGGAGA GUACCCACAU ACUCUGAUGA UCCUUCGGGA UCAUUCAUGG CAA |
| 5S-control | LR04 | GGGCCUUCGG GCCAAGUCUA CGGCCAUACC ACCCUGAACG CGCCCGAUCU CGUCUGAUCU CGGAAGCUAA GCAGGGUCGG GCCUGGUUAG UACUUGGAUG GGAGACCGCC UGGGAAUACC GGGUGCUGUA GGCGUCGACU UGCCAUGUGU AUGUGGGGAA ACCCACAUAC UCUGAUGAUC CUUCGGGAUC AUUCAUGGCA A |
| 5S-Mango with sequencing adapters | LR05 | GGGCUACACG ACGCUCUUCC GAUCUGUCUA CGGCCAUACC ACCCUGAACG CGCCCGAUCU CGUCUGAUCU CGGAAGCUAA GCAGGGUCGG GCCUGGUUAG UACUUGGAUG GGAGACCGCC UGGGAAUACC GGGUGCUGUA GGCGUCGACU UGCCAUGUGU AUGUGGGUAC GAAGGAGAGG AGAGGAAGAG GAGAGUACCC ACAUACUCUG AUGAUCCUUC GGGAUCAUUC AUGGCAAAGA UCGGAAGAGC ACACGUCU |
| Mango construct for mutational profiling | LR06 | GGGCUACACG ACGCUCUUCC GAUCUCUGGG UCGCAGUAAC CCCAGUUAAC AAAACAAGGU UCGACUUGCC AUGUGUAUGU GUGGUACGAA GGAGAGGAGA GGAAGAGGAG AGUACCCAUC AUACUCUGAU UGAUCCUUCG UGGAUCAUUC AUUGGCAAAG AUCGGAAGAG CACACGUCU |
| 5S binder | FZ01 | GGAACCUUCG ACGCCUACAG CACCCGGUAU UCCCAGGCGG UCUCCCAUCC AAGUA |

##### 1.2 DNA sequences

| Name | ID | Sequence |
|---|---|---|

| | | |
|---|---|---|
| 5S-Mango template (amplified from gBlocks Gene Fragments from IDT) | LR03D | GCGGAATTCT AATACGACTC ACTATAGGTC TACGGCCATA CCACCCTGAA CGCGCCCGAT CTCGTCTGAT CTCGGAAGCT AAGCAGGGTC GGGCCTGGTT AGTACTTGGA TGGGAGACCG CCTGGGAATA CCGGGTGCTG TAGGCGTCGA CTTGCCATGT GTATGTGGGT ACGAAGGAGA GGAGAGGAAG AGGAGAGTAC CCACATACTC TGATGATCCT TCGGGATCAT TCATGGCAA |
| 5S-control template (amplified from gBlocks Gene Fragments from IDT) | LR04D | GCGGAATTCT AATACGACTC ACTATAGGGC CTTCGGGCCA AGTCTACGGC CATACCACCC TGAACGCGCC CGATCTCGTC TGATCTCGGA AGCTAAGCAG GGTCGGGCCT GGTTAGTACT TGGATGGGAG ACCGCCTGGG AATACCGGGT GCTGTAGGCG TCGACTTGCC ATGTGTATGT GGGGAAACCC ACATACTCTG ATGATCCTTC GGGATCATTC ATGGCAA |
| Template for 5S-Mango with sequencing adapters (amplified from gBlocks Gene Fragments from IDT) | LR05D | TAATACGACT CACTATAGGG CTACACGACG CTCTTCCGAT CTGTCTACGG CCATACCACC CTGAACGCGC CCGATCTCGT CTGATCTCGG AAGCTAAGCA GGGTCGGGCC TGGTTAGTAC TTGGATGGGA GACCGCCTGG GAATACCGGG TGCTGTAGGC GTCGACTTGC CATGTGTATG TGGGTACGAA GGAGAGGAGA GGAAGAGGAG AGTACCCACA TACTCTGATG ATCCTTCGGG ATCATTCATG GCAAAGATCG GAAGAGCACA CGTCT |
| Template for Mango construct for mutational profiling (ordered as Ultramer DNA from IDT) | LR06D | AGACGTGTGC TCTTCCGATC TTTGCCAATG AATGATCCAC GAAGGATCAA TCAGAGTATG ATGGGTACTC TCCTCTTCCT CTCCTCTCCT TCGTACCACA CATACACATG GCAAGTCGAA CCTTGTTTTG TTAACTGGGG TTACTGCGAC CCAGAGATCG GAAGAGCGTC GTGTAGCCCT ATAGTGAGTC GTATTA |
| 5S binder template (amplified from gBlocks Gene Fragments from IDT) | FZ01D | GCGGAATTCT AATACGACTC ACTATAGGAA CCTTCGACGC CTACAGCACC CGGTATTCCC AGGCGGTCTC CCATCCAAGT A |
| T7 promotor (for IVT of LR06) | LR06Pr | TAATACGACT CACTATAGGG |

##### 1.3 Primers

| Name | ID | Sequence |
|---|---|---|
| 5S-M/C Fwd primer | LR034F | GCGGAATTCT AATACGACTC ACTATA |
| 5S-M/C Rev primer | LR034R | TTGCCATGAA TGATCCCGAA |

| | | |
|---|---|---|
| SeqAdapt Fwd primer | LR05F | TAATACGACT CACTATAGGG CTACACGA |
| SeqAdapt Rev primer (also used as RT primer) | LR05R | AGACGTGTGC TCTTCCGATC T |
| F-in (RT-qPCR) | WW01f | GAATACCGGG TGCTGTAGGC |
| R-in (RT-qPCR) | WW01r | AGACGTGTGC TCTTCCGATC T |
| F-ex (RT-qPCR) | WW02f | CTACACGACG CTCTTCCGAT CT |
| R-ex (RT-qPCR) | WW02r | CCTACAGCAC CCGGTATTCC |

#### 2. Custom light chamber setup

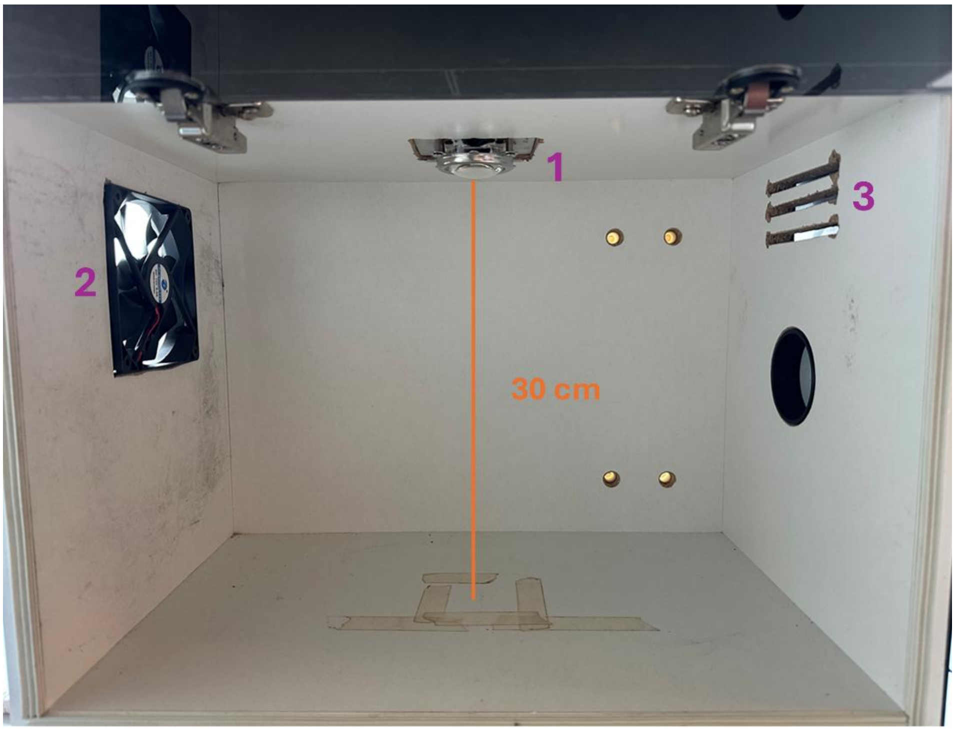


1. Chip-on-board LED (Chanzon, DC 30V - 34V / 100 Watt), equipped with a 120-degree lens, a heat sink and a fan.
2. Cooling fan from the side.
3. Ventilation slots.

Samples are placed at the center of the bottom, approximately 30 cm away from the CoB.

#### 3. In vitro transcription (IVT)

##### 3.1 DNA Templates

5S-Mango (LR03D), 5S-control (LR04D), and sequencing adapter-containing Mango DNA (LR05D) templates obtained from IDT were amplified using Phusion™ High-Fidelity DNA Polymerase. Briefly, DNA template (20 ng) was combined with the corresponding forward and reverse primers (1 µM each), Phusion HF PCR buffer (1x, NEB #B0518S), dNTPs (0.2 mM each), and Phusion DNA polymerase (0.02 U/µL, ThermoFisher F-530L) in a total reaction volume of 100 µL. Specifically, primers LR034F and R were for amplifying LR03D and LR04D; primers LR05F and R were for LR05D. PCR amplification was performed with an initial denaturation step at 98 °C for 30 s, followed by 30 cycles of denaturation at 98 °C for 10 s, annealing at 58 °C for 30 s, and extension at 72 °C for 30 s, with a final extension at 72 °C for 5 minutes. PCR products were purified using a MinElute PCR Purification Kit, and DNA concentration was determined by NanoDrop spectrophotometry.

For Mango used in mutational profiling, the template LR06D was ordered as single-stranded Ultramer DNA, which is reverse complementary to the IVT RNA sequence. LR06D was annealed with a T7 promoter oligo LR06Pr by combining both oligonucleotides at 20 µM in a total volume of 10 µL nuclease-free water, followed by heating to 95 °C for 3 minutes and gradually cooling to 20 °C over 30 minutes. The resulting annealed template was used directly for IVT.

##### 3.2 Small-scale IVT

DNA template (1 µg) was transcribed in a 20 µL reaction containing T7 transcription buffer (0.75x, NEB B9012), DTT (5 mM), NTPs (2.5 mM each), SUPERase·In™ RNase Inhibitor (0.5 U/µL, Invitrogen AM2694), and recombinant T7 RNA polymerase produced in-house (0.25 µg/µL, p6XHis-T7(P266L). The plasmid was a gift from Anna Pyle (Addgene ID 174866). The transcription reaction was incubated at 37 °C for 16 hours. Residual DNA template was subsequently degraded by incubating with TURBO DNase™ (0.8 U/µL, Invitrogen AM2238) at 37 °C for 30 minutes.

RNA generated by small-scale IVT was purified with Dynabeads™ MyOne™ Silane beads. Briefly, 2 mg beads were washed three times with 300 µL of RLT buffer (Qiagen 79216). RNA was bound by combining the transcription mixture with 350 µL RLT buffer (Qiagen), 10 µL of 5 M NaCl, and 615 µL absolute ethanol, followed by gentle inversion at room temperature for 15 minutes. The beads were subsequently washed three times with 200 µL of 75% ethanol, air-dried, and the RNA was eluted in 20 µL of DEPC-treated water at room temperature for 1 minute. RNA concentration was determined by NanoDrop spectrophotometry.

##### 3.3 Large-scale IVT

The small-scale reaction conditions were scaled up to a final volume of 100 µL reaction and incubated at 37 °C for 4 hours, followed by degradation of residual DNA template with TURBO DNase™ at 37 °C for 30 minutes.

For large-scale IVT of the RNA for Next-Generation sequencing (LR06), the template DNA (LR06D) was ordered as single-stranded Ultramer from IDT. To perform IVT, a DNA oligo was annealed to the T7 promoter region for T7 RNA polymerase recognition. The annealed DNA template (0.4 µM) was transcribed in a 50 µL reaction containing T7 transcription buffer, DTT (5 mM), NTPs (7.5 mM each), $MgCl_2$ (30 mM), SUPERase·In™ RNase Inhibitor (0.5 U/µL), and recombinant T7 RNA polymerase produced in-house (0.25 µg/µL). The transcription reaction was incubated at 37 °C for 4 hours, followed by degradation of residual DNA template with TURBO DNase™ (0.8 U/µL) at 37 °C for 30 minutes.

RNA generated by large-scale IVT was purified by preparative PAGE. RNA was mixed with loading buffer containing formamide (47.5%) and EDTA (12.5 mM) in DEPC-treatet water, denatured at 95 °C for 3 minutes, and immediately cooled on ice for 5 minutes. Subsequently, 70 µL of the denatured transcription reaction was loaded per lane onto a preparative 6% denaturing urea PAGE gel (19:1 acrylamide/bisacrylamide, 1.5 mm thick, 14 cm x 15 cm) and subjected to electrophoresis at 250 V for 110 minutes using a Hoefer SE400 system. Product bands were identified by UV shadowing at 254 nm (1 – 2s exposure) on a TLC plate, excised, transferred to tubes, and crushed. The crushed gel was soaked in 500 µL of elution buffer (500 mM $NH_4OAc$, 1 mM EDTA) and the slurry was gently rotated at room temperature for 1 hour. The RNA was recovered by centrifugation at 15 000 x g for 5 minutes and collecting the supernatant. The supernatant was further filtered through a 0.2 µm PTFE filter and further purified using Zymo-Spin IIICG Columns according to manufacturer instructions. Purified RNA was eluted in 50 µL per column. The gel slurry above can be stored at least overnight at -20 °C.

#### 4. General procedures for RNA labeling, purification and analysis

##### 4.1 Small molecule-directed proximity labeling

RNAs (0.5 µM) were folded in 1x folding buffer (10 mM HEPES, adjusted to pH 7.5 with 5.25 mM KOH, 135 mM KCl in DEPC-treated water) by heating to 90 °C for 5 minutes, followed by cooling to 20 °C at a rate of 1 °C $s^{-1}$. To induce proximity-dependent labeling, **TO-EY** was added, and the mixture was incubated for 30 minutes at room temperature. Then **AnA** or **APA** probe was added (0.1 mM), followed by irradiation with green or blue light for 10 minutes in a custom-built light chamber (see **Section 2** for the setup), unless otherwise mentioned.

##### 4.2 Post-labeling clean-up and analysis

The labeled RNA was immediately subjected to the extraction using Acid Guanidinium thiocyanate-Phenol-Chloroform (AGPC), followed by purification with the RNA Clean & Concentrator™-5 kit (Zymo Research) following the supplier's standard protocol.

For in-gel fluorescence detection, Cy5 was conjugated to labeled RNA via strain-promoted azide-alkyne cycloaddition (SPAAC). Briefly, RNA was incubated with DBCO-Cy5 (250 µM, Lumiprobe cat. 433F0) in formamide (47.5%) containing EDTA

(12.5 mM) at 55 °C for 10 minutes. Cy5 labeled RNA was subsequently purified using the same AGPC extraction and clean-up procedure as described above. For the RNAse-treated samples, 1 µL RNAse cocktail (10 U/mL RNAse A and 400 U/mL RNAse T1, Invitrogen AM2286) was added after the second RNA purification step, followed by incubation at room temperature for 15 minutes. The digested samples were analyzed directly without further purification.

For analysis, RNA samples were denatured in 6 M urea by heating to 95 °C for 3 minutes, followed by immediate cooling on ice for 5 minutes. Samples were resolved on a 6% or 8% denaturing polyacrylamide gel (29:1 acrylamide:bisacrylamide, urea-TBE) at 250 V for 30 minutes. Gels were imaged for Cy5 fluorescence using a Bio-Rad ChemiDoc™ MP imaging system. To verify comparable RNA loading, gels were subsequently stained with ethidium bromide (EtBr) (40 µg in 40 mL of 1x TBE) and reimaged to assess the total RNA loading.

Cy5 fluorescence band intensities were quantified as background-corrected integrated volumes using Bio-Rad Image Lab software (v6.1, Bio-Rad Laboratories). The band volumes were corrected for slight differences in RNA loading by scaling each value to the corresponding EtBr RNA loading signal relative to the mean RNA loading signal.

#### 5. Experiment-specific labeling procedures

##### 5.1 Efficiency comparison between alkyl amine and aniline

The procedure corresponds to **Figure 2C**, **D**, **E** and **S1**. 5S-control RNA (LR04) was used to evaluate the labeling efficiency of the **APA** and **AnA** probes. 5S-control RNA was generated following the small-scale IVT protocol with LR04D as the templatee. Labeling, clean-up and analysis were performed as described in the general labeling procedure above without folding and small molecule binding, using free EY (10 µM) as photosensitizer and **APA** or **AnA** as probe.

##### 5.2 Time-dependent RNA labeling

The procedure corresponds to **Figure S2**. 5S-control RNA (LR04) was used to check the dependency on time of light exposure for the **APA** and **AnA** probes (0.1 mM). 5S-control RNA was generated by large-scale IVT using LR04D as template. Labeling was performed using free EY (10 µM) as photosensitizer without RNA folding or ligand binding. Aliquots (10 µL each) of the labeling mixture in different tubes were irradiated simultaneously, and individual aliquots were collected at 1-minute intervals over a total irradiation period of 11 minutes. AGPC was immediately added to each tube once taken from the chamber and placed in dark on ice for further processing until all samples were collected for batch purification.

##### 5.3 Selectivity comparison between alkyl amine and aniline

The procedure corresponds to **Figure 3D**, **E** and **S3**. 5S-Mango (LR03) and 5S-control (LR04) RNA were used to evaluate the labeling selectivity, either individually or in equimolar mixed samples. RNAs were generated by large-scale IVT with the templates LR03D and LR04D, respectively, followed by purification by preparative PAGE. Labeling, clean-up and analysis were performed according to the general procedure described above, using **TO-EY** (5 µM) and **APA** or **AnA** (0.1 mM) as probe.

##### 5.4 Effect of Mango-binding competitor on labeling selectivity

The procedure corresponds to **Figure S4**. 5S-Mango (LR03) and 5S-control (LR04) RNA were mixed at equimolar concentrations to assess the effect of competitive binding on labeling selectivity. Both RNA constructs were generated by large-scale IVT (LR03D and LR04D, respectively). Labeling, clean-up and analysis were performed according to the general labeling procedure using **AnA** (0.1 mM) as probe. Competition was introduced by incubating the RNA with **TO-EY** (10 µM) as photosensitizer and **TO-amine** at concentrations ranging from 5-200 µM as competitor during RNA folding.

##### 5.5 Effect of TO-EY concentration on labeling selectivity

The procedure corresponds to **Figure 3F** and **G**. 5S-Mango (LR03) and 5S-control (LR04) RNAs were used in equimolar mixed samples to evaluate the effect of **TO-EY** concentration on labeling selectivity. RNAs were generated from large-scale IVT using LR03D and LR04D as DNA template, respectively. Labeling, clean-up and analysis were performed according to the general labeling procedure described above, using **TO-EY** concentration ranging from 0.1 to 100 µM and **AnA** (0.1 mM) as probe.

##### 5.6 Effect of Singlet Oxygen quenching on labeling selectivity

The procedure corresponds to **Figure 3H** and **I.** 5S-Mango (LR03) and 5S-control (LR04) RNA were used in equimolar mixed samples to evaluate the effect of sodium azide ($NaN_3$) on labeling selectivity. RNAs were generated from large-scale IVT using LR03D and LR04D as DNA template, respectively. Labeling, clean-up and analysis were performed according to the general procedure described above in the presence of 10 µM **TO-EY**, $NaN_3$ concentrations ranging from 0 to 50 mM, and 0.1 mM **AnA** as probe. For this reaction, a commercial light chamber (PhotoRedOx Box, 525 nm LED, HaptoChem) was used.

##### 5.7 Proximity labeling of 5S binder

The procedure corresponds to **Figure S5**. 5S binder RNA (FZ01) was hybridized to either 5S-Mango (LR03) or 5S-control (LR04) RNA. All RNA constructs were generated by large-scale IVT from the DNA templates FZ01D, LR03D and LR04D, respectively. 5S binder RNA was annealed to either 5S-Mango or 5S-control RNA at equimolar concentrations in 1x folding buffer by heating to 95 °C for 3 minutes, followed by gradual cooling to 20 °C over 30 minutes. The general labeling, clean-up and analysis procedure were subsequently performed on the resulting RNA duplex (2 µM final concentration) with either 10 µM **TO-EY** or free EY, and with **AnA** as the probe. Labeled products were analyzed on by 8% denaturing PAGE (29:1 acrylamide:bisacrylamide).

##### 5.8 Proximity of RNA for pairwise RT stop and qPCR

The procedure corresponds to **Figure 4B**. Sequencing adapter-containing 5S-Mango RNA construct (LR05) was generated by large-scale IVT from the template LR05D, followed by purification by preparative PAGE. Labeling was performed according to the general labeling procedure described above, using 10 μM **TO-EY**, 10 mM $NaN_3$ and LR05 RNA at a concentration of 1.2 μM in the HaptoChem PhotoRedOx Box. Samples were purified following the general clean-up procedure.

##### 5.9 Proximity labeling of RNA for mutational profiling (MaP)

The procedure corresponds to **Figure 5**. Mango construct for NGS (LR06) was generated by IVT with the single-stranded DNA template LR06D annealed to a T7 promoter DNA oligo LR06Pr. Labeling was performed according to the general labeling procedure described above with **TO-EY** (10 μM) in the presence of $NaN_3$ (5 mM), using **AnA** (0.1 mM) as probe, in the HaptoChem PhotoRedOx Box. Samples were purified following the general clean-up procedure.

#### 6. Other procedures

##### 6.1 Mango RNA binding experiment with fluorescence readout.

The protocol corresponds to **Figure 3B**. Mango-II RNA aptamer LR01 (0.1 μM) was used to evaluate the binding of **TO-amine** (5 μM). A non-binding RNA oligonucleotide LR02 was included as a negative control. Both RNA oligonucleotides were obtained from Integrated DNA Technologies (IDT). Folded RNA and **TO-amine** were mixed in 1x folding buffer in a black 384-well microplate and incubated at room temperature in the dark for 30 minutes. Fluorescence intensity was measured using a BMG Labtech POLARstar Omega plate reader, with excitation at 485 nm and emission detection at 520 nm.

##### 6.2 Pairwise RT stop assay and qPCR analysis in vitro and in cell

For in-vitro labeled RNA, the RT-qPCR was performed after labeling and clean-up. The RNA sample per condition was diluted to 20 ng/mL with nuclease-free water and equally split into two (one for Mango-excluding RT and the other for Mango-including RT). PCR reactions were set up following manufacturer's instruction (Luna Universal One-Step RT-qPCR Kit, E3005S, NEB) and was performed in 3 technical replicates. For each reaction, 5 μL reaction mix, 0.5 μL WarmStart RT Enzyme Mix, 0.4 μL primers (WW01f/r or WW02f/r) and 3.1 μL nuclease-free water were mixed in a 200 μL PCR tube (total volume 9 μL). To this mixture was added 1 μL diluted RNA solution. The qPCR was performed using a BioRad CFX96 PCR Detection System following kit manufacturer's instruction.

For in-cell proximity labeling, HEK293T cells were seeded into 12-well plates and culture to 80% confluency prior to transfection. To prepare pre-complexed Mango-TO-EY for transfection, 1 μg folded Mango RNA (LR05) and was added into 50 μL Mango binding buffer (140 mM KCl, 1 mM $MgCl_2$ in 1x DPBS). In a new tube, 2 μL

Lipofectamine RNAiMAX was added into serum-free DMEM supplemented with 140 mM KCl and 1 mM $MgCl_2$. Mix the solutions in the two tubes at 21 °C for 15 min. The old culture media was replaced with the above transfection mix in serum-free DMEM supplemented with 140 mM KCl and 1 mM $MgCl_2$. The cells were incubated for 3 hours, after which the cells were washed once with 0.5 mL Mango binding buffer. A solution of biotin-$PEG_4$-aniline (250 μM) in 0.5 mL Mango binding buffer was added to the cells, which is kept in dark for 5 min. The cells were irradiated under green light for 5 min. The cells were then washed twice with Mango binding buffer and lysed in TRIzol. Total RNA was extracted following the supplier's protocol.

For each RNA sample, 10 μL MyONE C1 streptavidin beads (Invitrogen, 35002D) were used. To prepare beads for binding, the storage solution was removed, and the beads were washed three times with 500 μL of bind and wash (B&W) buffer (10 mM Tris-HCl, pH 7.5; 1 mM EDTA; 2M NaCl; 0.1% Tween-20). The beads were taken up in 500 μL new B&W buffer, into which 2 μg extracted total RNA was added. The slurry was incubated at 4 °C for 1 hour. The RNA bound beads were washed 3 times with B&W buffer. After removing the last wash, 100 μL 1x RT buffer (Thermo Scientific, 00758185) was added to each sample. The slurry was thoroughly mixed before being split into two new tubes (one for F-ex/R-ex primers, the other for F-in/R-in primers).

RT reaction mix (Thermo Scientific, 18080044) was prepared as follows. The mix1 contains 0.4μL primers, 1 μL 10mM dNTP, and toppe up with nuclease-free water to a total of 12 μL. The mix2 contains 4 μL 5x First-Strand Buffer, 1 μL 0.1 M DTT, 0.5 μL RNase Inhibitor, and 0.5 μL SuperScript™ III, and topped up nuclease-free water to 7 μL. First, the supernatant RT buffer was aspirated on magnet, and mix1 was added to the beads. The slurry was heated at 65 °C for 5 min and placed on ice for 1 min, to which the mix2 was then added. The complete RT reactions were incubated on a thermoshaker at 55 °C, 400 rpm for 1 hour. The slurry was then heated up to 95 °C for 5 min and immediately placed on the magnet for transferring the cDNA-containing supernatant to a new tube.

The qPCR reaction solution master mix was prepared according to manufacturer's guide (MedChemExpress, HY-K0501). Each 200 μL PCR tube contains 8 μL reaction solution, 5 μL Sybr master mix, 0.4 μL primer pair (10 μM), and was topped up to 18 μL with nuclease-free water. 2 μL cDNA solution from above was added to the tube. Real-time qPCR was performed using a CFX96 real-time system (BioRad, C1000 Touch Thermal Cycler).

For quantification, Ct values from each individual primer pair were normalized over the mean of Mango only (unmodified) control of the same primer pair (by subtraction). The apparent quantity of RT products was calculated as follows

$$\text{Apparent quantity} = 2\text{\^{}}[-(Ct_{modified} - Ct_{unmodified})],$$

where the $Ct_{modified}$ is the normalized mean of 3 technical replicates of labeled samples (either randomly or proximity labeled); and $Ct_{unmodified}$ is the normalized mean of 3 technical replicates of unlabeled control.

##### 6.3 Confocal imaging studies

HEK293T cells were seeded into 8-well ibidi plates. Transfection was performed when the confluency reached 70%. 0.3 μg control or Mango RNA was incubated with 1 μM **TO-amine** in 50 uL Mango binding buffer for 10 min. In a new tube, 0.6 μg Lipofamine RNAiMAX was added in 50 μL DMEM with 140 mM KCl and 1 mM $MgCl_2$, which was incubated for 15 min. The above two solutions were mixed at equal volume. Culture medium was aspirated and replaced with 200 μL DMEM containing 140 mM KCl 1 mM $MgCl_2$. The 100 μL RNA-containing transfection solution above was added into the cells, which was incubated for 3 hours. After transfection, the cells were fixed with 4% PFA on ice for 10 min. The fixed cells were washed 3 times with Mango binding buffer, and permeabilized using 0.2% Triton X-100 and on ice for 10 min. The cells were then washed twice with Mango binding buffer. Nucleus was stained with 1 μg/mL DAPI for 5 min. The cells were then washed twice with Mango binding buffer and taken to the microscope for imaging. DAPI, excitation 405 nm, filter 450 – 470 nm; Mango fluorescence, excitation 488 nm, filter 510 – 555 nm.

##### 6.4 Next-Generation sequencing library preparation

Labeled and purified RNA (100 ng) was combined with the RT primer (1 μM) and dNTPs (0.5 mM), heated to 70 °C for 5 minutes, and immediately cooled on ice for 2 minutes. Reverse transcription was initiated by addition of Tris-HCl buffer (50 mM, pH 8), KCl (75 mM), DTT (10 mM), $MnCl_2$ (6 mM), SUPERase·In™ RNase Inhibitor (0.5 U/μL), and SuperScript™ II (10 U/μL, Invitrogen 18064014) in a total volume of 10 μL. Reactions were incubated at 42 °C for 90 minutes, followed by sequential incubation at 50 °C, 55 °C, and 60 °C for 10 minutes each, and a final incubation at 75 °C for 15 minutes.

Resulting cDNA was purified using the Zymo RNA Clean & Concentrator™-5 columns. Briefly, 20 μL binding buffer and 50 μL ethanol absolute were added to the cDNA sample, which was then transferred to the spin column and centrifuged at 16 000 x g for 30 s. Columns were equilibrated with 400 μL RNA prep buffer and centrifugation at 16 000 x g for 30 s, followed by washing with 700 μL RNA wash buffer and centrifugation at 16 000 x g for 30 s, and a second wash with 400 μL RNA wash buffer, followed by centrifugation at 16 000 x g for 1 minute. Purified RT product was eluted in 10 μL nuclease-free water following incubation at room temperature for 2 minutes and centrifugation at 16 000 x g for 1 minute.

Libraries were then amplified using NEBNext® Ultra™ II Q5® Master Mix (NEB, M0544L) and barcoded primers, as per manufacturer instructions.

##### 6.5 Sequencing data analysis

Read mapping was performed using the rf-map module of the RNA Framework[1] and Bowtie2 v2.3.5.1[2] (parameters: -b2 -ctn -cmn 0 -mp "--very-sensitive-local" -bnr). Mutations were then counted using the rf-count module (parameters: -m -rd).

Mutation rates for each nucleotide were log-transformed and analyzed using the limma package (v3.66.0)[3] with empirical Bayes variance moderation. Pairwise comparisons were performed between **TO-EY** labeling and free EY labeling. Red bars in the graph indicate nucleotides with a positive log fold change and an adjusted p-value < 0.05 (Benjamini–Hochberg false discovery rate correction).

**6.6 Statistical Analysis**

One-way ANOVA with Šidák's multiple-comparisons correction was performed for **Figure 2C**, **D** and **S1** following log transformation of data. Unpaired t-test was performed for **Figure 2E**. One-way ANOVA with Dunnett's multiple-comparisons test was performed for **Figure 3B**. Two-way ANOVA with Tukey's multiple-comparisons test was performed for **Figure 3E** and **S3**. limma differential analysis with Benjamini–Hochberg correction was performed for **Figure 5D**.

#### 7. Synthesis procedures

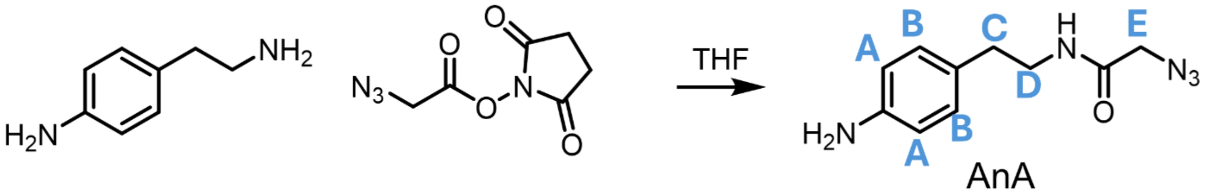


N-(4-aminophenethyl)-2-azidoacetamide (**AnA**). In a 25 mL round bottom flask, 2-azidoacetic acid NHS ester (25 mg, 0.13 mmol, 1.0 eq.) was dissolved in anhydrous 2 mL THF. 4-(2-aminoethyl)aniline (18 mg, 0.13 mmol, 1.05 eq.) was added to the solution. The reaction was stirred at 21 ℃ for 60 min. The solution was concentrated *in vacuo* and purified using silica gel chromatography. Yield, 87%.$^1$H-NMR (400 MHz, $CD_3OH$) δ 7.00 (d, J = 8.3 Hz, 2H, H-B), 6.71 (d, J = 8.3 Hz, 2H, H-A), 3.86 (s, 2H, H-E), 3.40 (t, J = 7.3 Hz, 2H, H-D), 2.71 (t, J = 7.3 Hz, 2H, H-C). HRMS (ESI, positive mode), $C_{10}H_{13}N_5O + H^+$, expected, 220.1198; observed, 220.1206.

$^1$H-NMR **AnA**

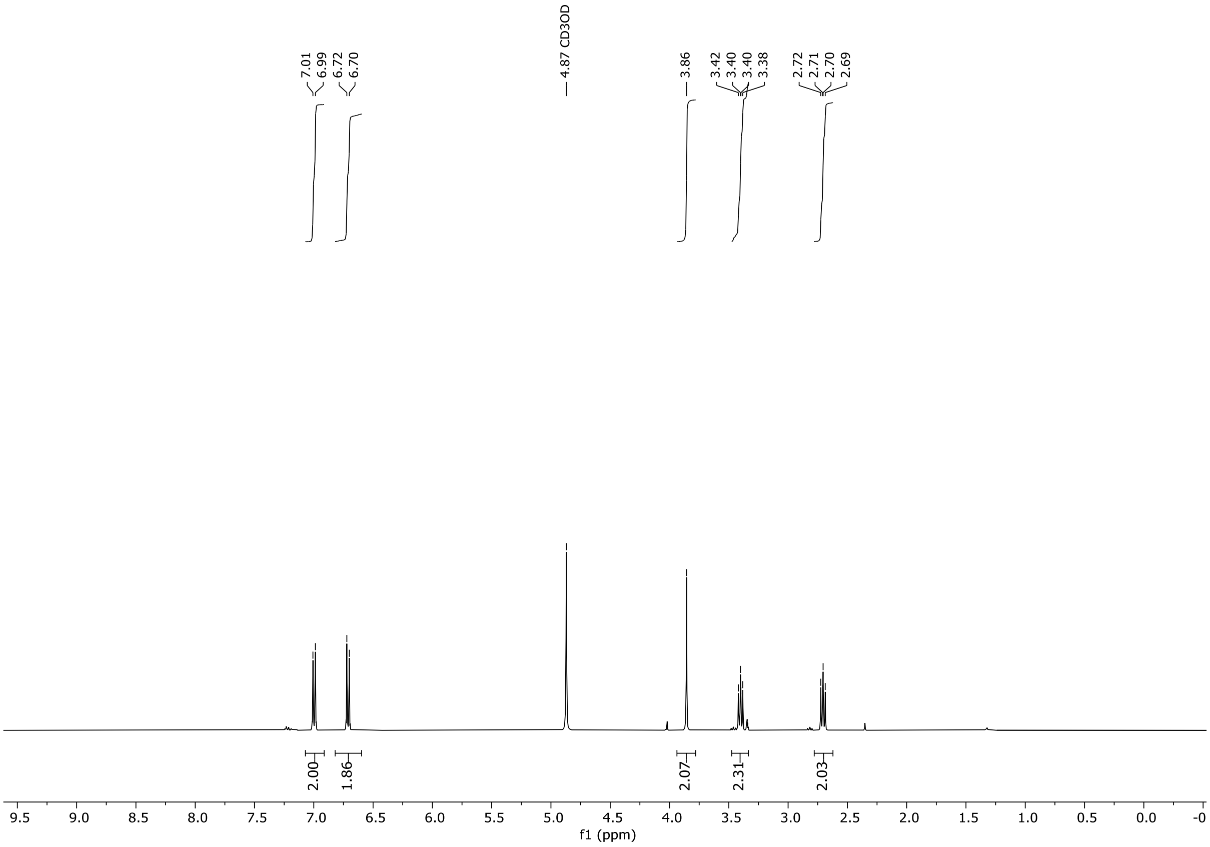

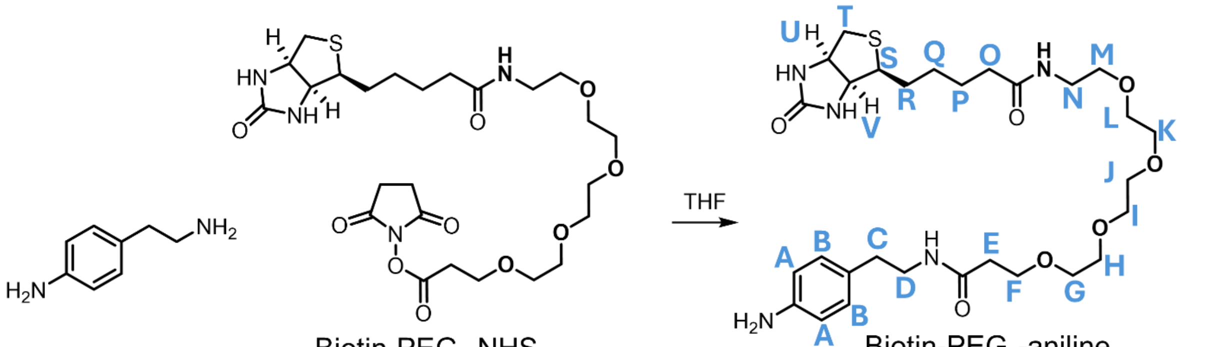


N-(4-aminophenethyl)-1-(5-((3aS,4S,6aR)-2-oxohexahydro-1H-thieno[3,4-d]imidazol-4-yl)pentanamido)-3,6,9,12-tetraoxapentadecan-15-amide (**biotin-PEG$_4$-aniline**). In a 15 mL falcon tube, Biotin-PEG$_4$-NHS (24 mg, 0.041 mmol, 1.0 eq.) was dissolved in anhydrous 2 mL THF. 4-(2-aminoethyl)aniline (7 mg, 0.049 mmol, 1.2 eq.) was added to the solution. The reaction was stirred at 21 ℃ for 60 min. The solution was concentrated *in vacuo*. The product was purified using silica gel chromatography. Yield, 21 mg, 84.7 %. $^1$H-NMR (400 MHz, $CDCl_3$) δ 7.00 – 6.96 (m, 2H, H-B), 6.93 (t, *J* = 5.5 Hz, 1H, N*H*), 6.68 – 6.60 (m, 2H, H-A), 6.56 (m, 1H, NH), 6.21 (s, 1H, N*H*), 5.38 (s, 1H, N*H*), 4.48 (dd, *J* = 7.9, 4.9 Hz, 1H, H-V), 4.29 (ddd, *J* = 7.7, 5.0, 2.5 Hz, 1H, H-U), 3.68 (t, *J* = 5.8 Hz, 2H, H-F), 3.65 – 3.58 (m, 8H, H-G/H/I/J/K/L/M), 3.58 – 3.49 (m, 6H, H-G/H/I/J/K/L/M), 3.47 – 3.36 (m, 4H, H-D/N), 3.12 (td, *J* = 7.3, 4.5 Hz, 1H, H-S), 2.89 (dd, *J* = 12.8, 4.9 Hz, 1H, H-T), 2.76 – 2.64 (m, 4H, H-T, H-C), 2.43 (t, *J* = 5.8 Hz, 2H, H-E), 2.18 (t, *J* = 7.4 Hz, 2H, H-O), 1.66 (dtq, *J* = 22.1, 14.8, 7.4 Hz, 4H, H-P/Q/R), 1.40 (p, *J* = 7.7 Hz, 2H, H-P/Q/R). HRMS (ESI, positive mode), $C_{29}H_{47}N_5O_7S + H^+$, expected, 610.3274; observed, 610.3270.

#### $^1$H-NMR **biotin-PEG$_4$-aniline**

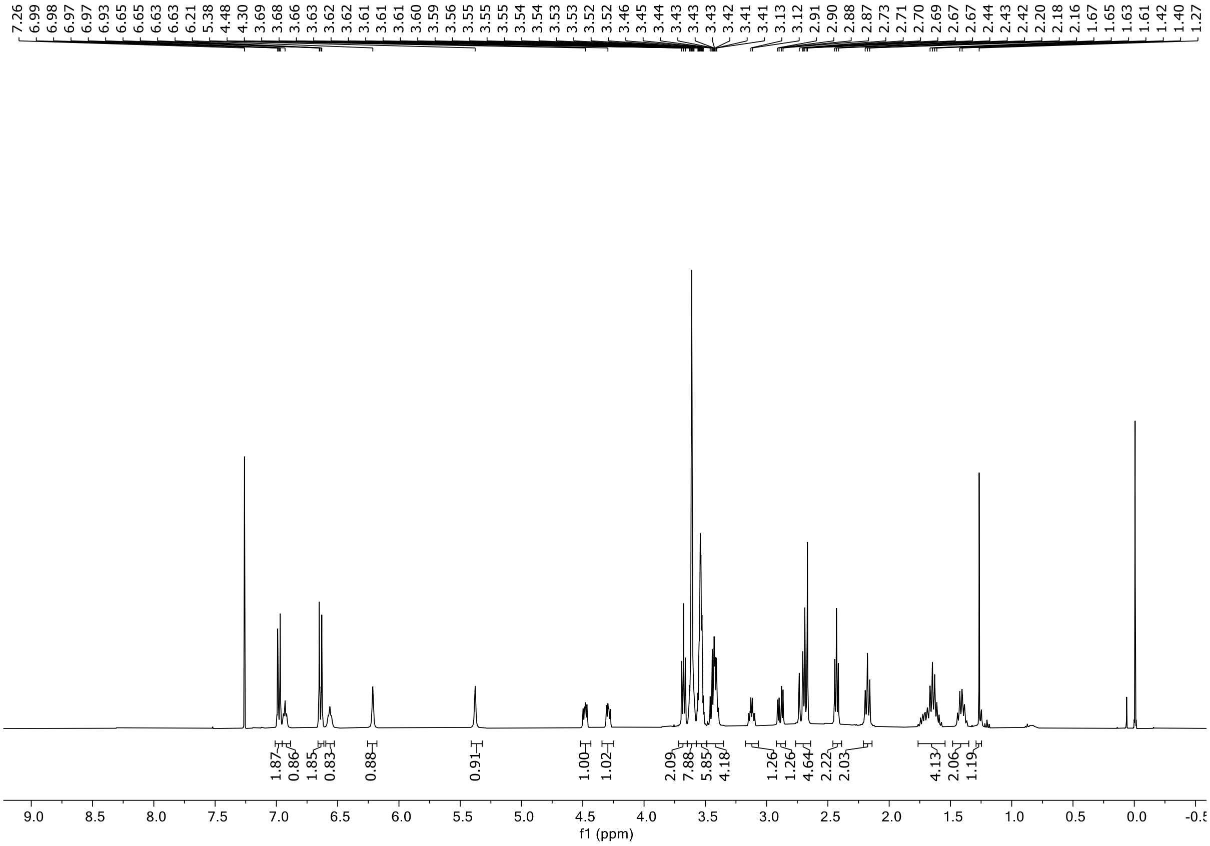


#### $^1$H-$^{13}$C-HSQC **biotin-PEG$_4$-aniline**

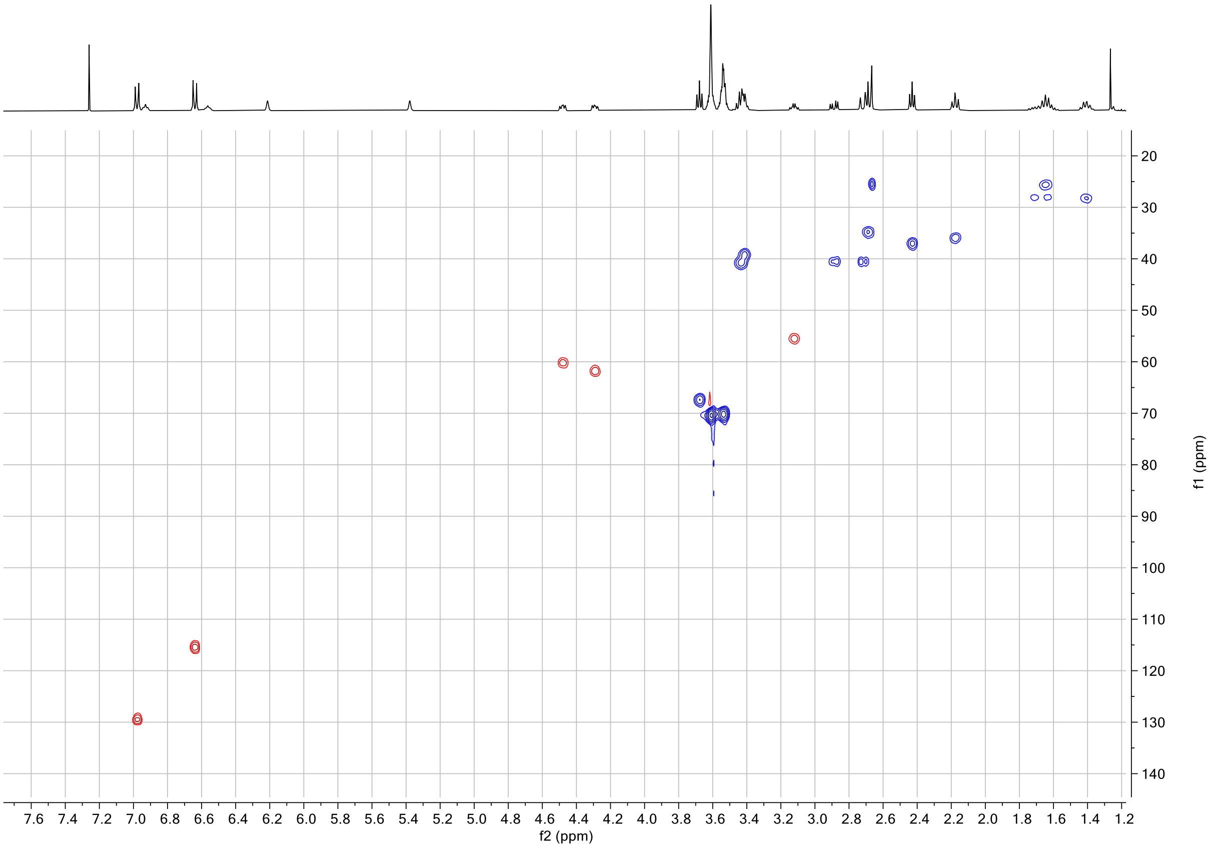

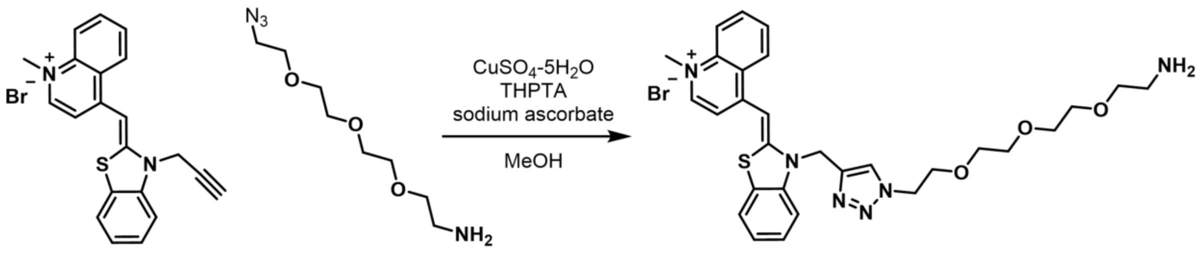


(Z)-4-((3-((1-(2-(2-(2-(2-aminoethoxy)ethoxy)ethoxy)ethyl)-1H-1,2,3-triazol-4-yl)methyl)benzo[d]thiazol-2(3H)-ylidene)methyl)-1-methylquinolin-1-ium bromide (**TO-amine**). TO-alkyne was obtained via reported procedure.[4] TO-alkyne (20 mg, 0.048 mmol, 1.0 eq.) and azide-PEG-amine (11 mg, 0.048 mmol, 1.0 eq) was dissolved in 1 mL MeOH in a Falcon tube. Premix $CuSO_4 \cdot 5H_2O$ (0.014 mmol, 0.3 eq., diluted from a 1 M stock solution), THPTA (0.029 mmol, 0.6 eq., diluted from a 0.1 M stock solution) in 200 µL milliQ. To the copper-ligand solution was added sodium ascorbate (0.072 mmol, 1.5 eq., diluted from a 0.5 M stock solution). Upon complete disappearance of the blue color, the copper solution was added to the azide and alkyne solution in MeOH. The reaction was vortexed for 10s at max speed and then place on a shaken at 21 ℃ for 1 hr in dark. Then 0.8 mL milliQ was added, and the mixture was spun down at 3000 g for 5 min. The supernatant was injected to a preparative HPLC system eluted with a 5 – 80% acetonitrile (in milliQ) gradient. Fractions containing the product was collected and lyophilized to afford **TO-amine** as a red powder. Isolated yield, 39.1%. $^1$H-NMR (600 MHz, $CD_3CN$) δ 8.70 (d, $J$ = 8.6 Hz, 1H), 8.30 (d, $J$ = 7.1 Hz, 1H), 8.16 (s, 1H), 8.05 – 7.97 (m, 2H), 7.91 (dd, $J$ = 7.8, 1.2 Hz, 1H), 7.86 – 7.79 (m, 2H), 7.68 – 7.62 (m, 1H), 7.48 – 7.42 (m, 2H), 7.35 (s, 1H), 5.75 (s, 2H), 4.54 (t, $J$ = 5.1 Hz, 2H), 4.16 (s, 2H), 3.84 (q, $J$ = 5.4 Hz, 2H), 3.67 – 3.57 (m, 2H), 3.55 – 3.45 (m, 2H), 3.43 – 3.35 (m, 2H), 3.10 – 3.05 (m, 2H). Skewed baseline in $^1$H-NMR was a result of solvent signal suppression (δ 1.97 ppm for acetonitrile and 2.52 ppm for DMSO. method, convolution; selectivity, 32). HRMS (ESI, positive mode), $C_{29}H_{35}N_6O_3S^+$, expected, 547.2486; observed, 547.2493.

#### $^1$H-NMR **TO-amine**

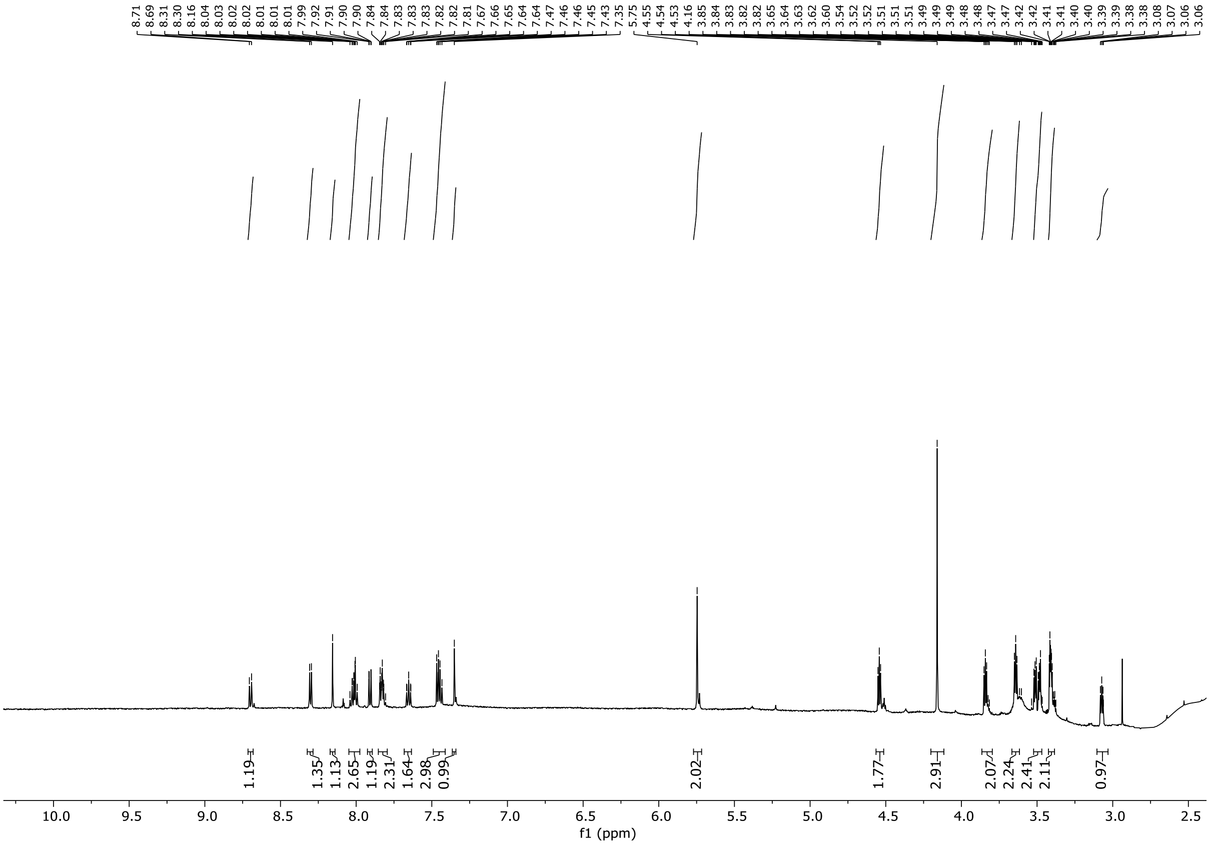


#### $^1$H-$^{13}$C-HSQC **TO-amine**

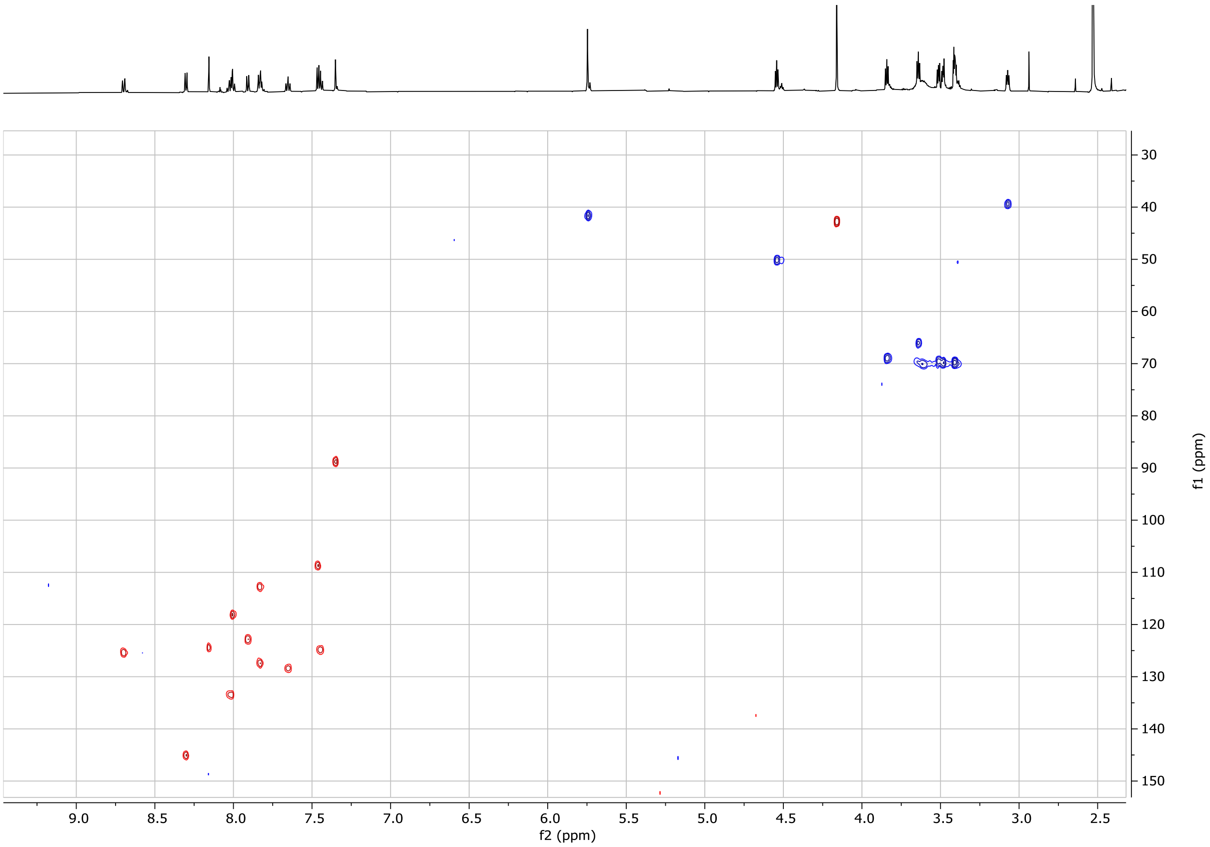

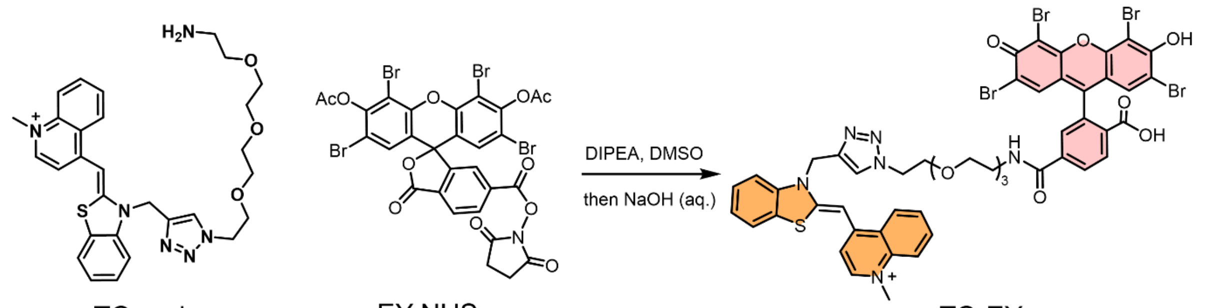


TO-amine EY-NHS TO-EY

(Z)-4-((3-((1-(1-(4-carboxy-3-(2,4,5,7-tetrabromo-6-hydroxy-3-oxo-3H-xanthen-9-yl)phenyl)-1-oxo-5,8,11-trioxa-2-azatridecan-13-yl)-1H-1,2,3-triazol-4-yl)methyl)benzo[d]thiazol-2(3H)-ylidene)methyl)-1-methylquinolin-1-ium bromide (**TO-EY**). In a 1.5 mL Eppendorf tube, TO-amine (2 mg, 3.2 μmol, 1.0 eq.) and DIPEA (0.5 μL, 3.2 μmol, 1.0 eq.) was dissolved in 400 μL DMSO. EY-NHS (1.1 mg, 1.3 μmol, 0.4 eq.), which was obtained via the reported procedure,[5] was dissolved in 50 μL DMSO, and added to the TO-amine solution. The reaction was shaken at 21 ℃ for16 hr. Then 2 μL NaOH solution (5 M stock solution, 10 μmol) was added to remove O-acetyl on EY. After 1 hr, the reaction was lyophilized. The product was purified via silica gel column chromatography to afford TO-EY as a magenta-red powder. Isolated yield, 57.2%. $^1$H-NMR (600 MHz, DMSO) δ 8.76 (d, J = 8.5 Hz, 1H), 8.64 – 8.57 (m, 2H), 8.30 (s, 1H), 8.05 – 7.91 (m, 6H), 7.78 – 7.72 (m, 1H), 7.60 – 7.55 (m, 2H), 7.37 (t, J = 7.6 Hz, 1H), 7.32 – 7.27 (m, 2H), 6.89 (s, 2H), 5.89 (s, 2H), 4.46 (t, J = 5.1 Hz, 2H), 4.17 (s, 3H), 3.70 (t, J = 5.1 Hz, 2H), 3.47 (t, J = 5.9 Hz, 2H). Skewed baseline in $^1$H-NMR was a result of solvent signal suppression (δ 2.52 ppm for DMSO. method, convolution; selectivity, 24). HRMS (ESI, positive mode), $C_{50}H_{41}Br_4N_6O_9S^+$, expected (highest relative abundance peak), 1220.9343; observed, 1220.9334.

#### $^1$H-NMR **TO-EY**

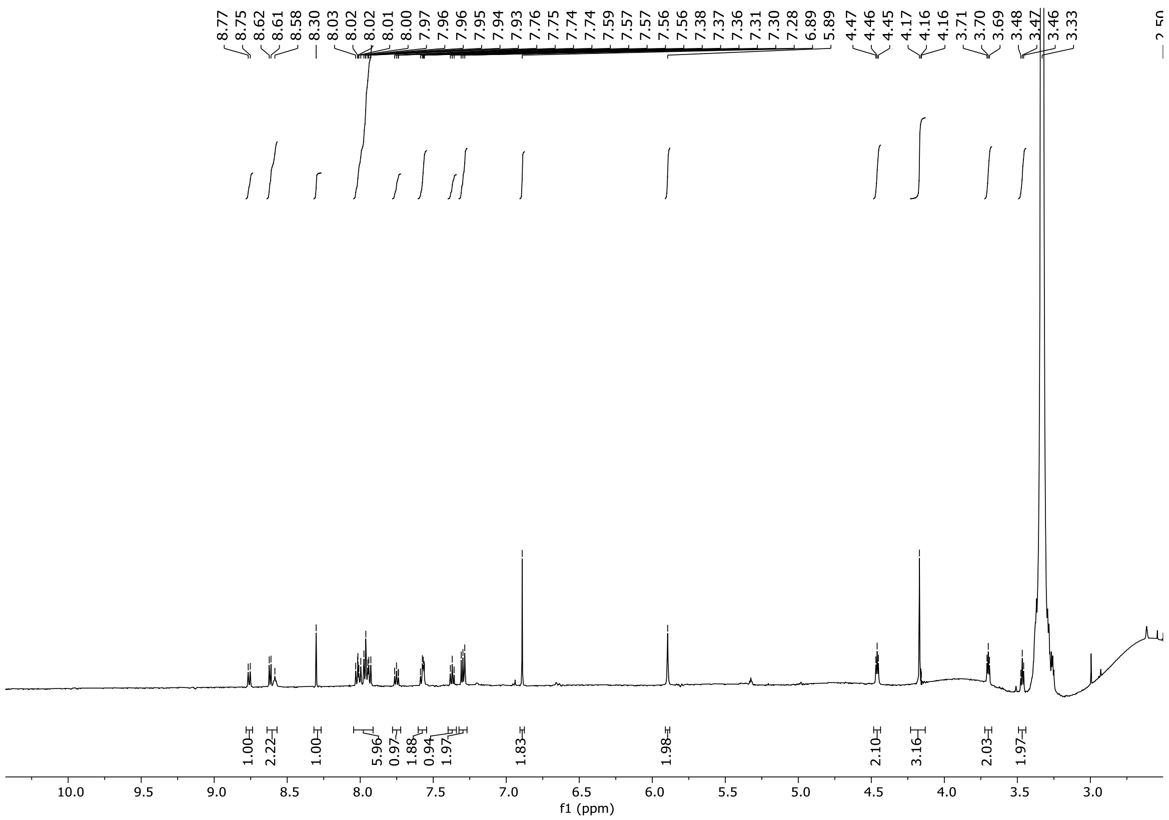

#### $^1$H-$^{13}$C-HSQC **TO-amine**

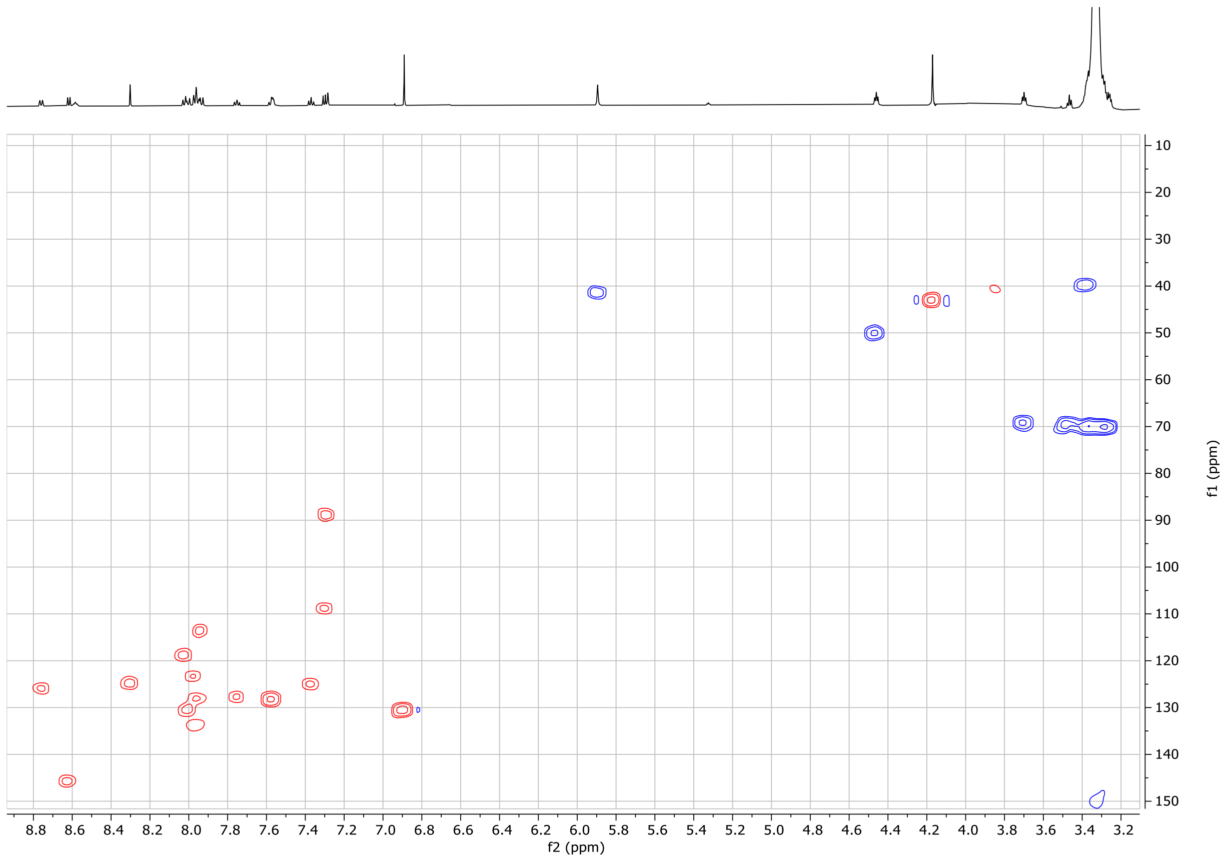

#### References


1. Incarnato, D., Morandi, E., Simon, L. M. & Oliviero, S. RNA Framework: an all-in-one toolkit for the analysis of RNA structures and post-transcriptional modifications. *Nucleic Acids Res* **46**, e97 (2018). https://doi.org/10.1093/nar/gky486
2. Langmead, B. & Salzberg, S. L. Fast gapped-read alignment with Bowtie 2. *Nat Methods* **9**, 357–359 (2012). https://doi.org/10.1038/nmeth.1923
3. Ritchie, M. E., Phipson, B., Wu, D. *et al.* limma powers differential expression analyses for RNA-sequencing and microarray studies. *Nucleic Acids Res* **43**, e47 (2015). https://doi.org/10.1093/nar/gkv007
4. Bychenko, O. S., Khrulev, A. A., Svetlova, J. I. *et al.* Red light-emitting short Mango-based system enables tracking a mycobacterial small noncoding RNA in infected macrophages. *Nucleic Acids Res* **51**, 2586–2601 (2023). https://doi.org/10.1093/nar/gkad100
5. Luo, H., Tang, W., Liu, H. *et al.* Photocatalytic Chemical Crosslinking for Profiling RNA-Protein Interactions in Living Cells. *Angew Chem Int Ed Engl* **61**, e202202008 (2022). https://doi.org/10.1002/anie.202202008
